# Tmn blocks phage spread via plasmolysis and triggers synergistic defence responses

**DOI:** 10.64898/2026.03.01.708804

**Authors:** Yi Wu, Zhiying Zhang, Sofya K. Garushyants, Ruizhe Li, Regan Doherty, James A. Milton, Matthew J. Cooper, Yilmaz Emre Gençay, Triana Amen, Somenath Bakshi, Dinshaw J. Patel, Eugene V. Koonin, Franklin L. Nobrega

**Affiliations:** School of Biological Sciences, Faculty of Environmental and Life Sciences, University of Southampton, Southampton, UK; Structural Biology Program, Memorial Sloan Kettering Cancer Center, New York, NY, USA; Division of Intramural Research, National Library of Medicine, National Institutes of Health, Bethesda, MD, USA; Department of Engineering, University of Cambridge, Trumpington Street, Cambridge, UK; Biomedical Imaging Unit, Faculty of Medicine, University of Southampton, Southampton, UK; School of Ocean and Earth Science, Faculty of Environmental and Life Sciences, University of Southampton, Southampton, UK; SNIPR Biome, Copenhagen, Denmark

## Abstract

Membrane-associated phage defences remain poorly understood. Here we characterise Tmn, a YobI-family transmembrane P-loop NTPase that protects bacteria from phage infection by establishing a plasmolysis-associated antiviral state. Upon recognising phage T2 RIIB protein, Tmn enhances ATP turnover and selectively exports Mg^2+^, causing rapid cytoplasmic collapse that arrests phage replication without detectable membrane depolarisation or gross leakage of the cell content. Cryo-electron microscopy shows that Tmn assembles into a decameric membrane complex with extended cytosolic arms, an uncommon architecture among P-loop NTPases. The cytosolic arms, including a solenoid-like repeat domain, mediate trigger interaction and determine specificity. In addition to its primary defence function, Tmn-driven ATP collapse activates otherwise silent ATP-depletion-sensing defences, including Gabija and Septu type I, providing a mechanistic basis for synergy among defence systems and limiting secondary phage spread. These findings demonstrate that Tmn is a membrane-integrated sensor-effector that couples phage recognition to metabolic collapse and coordinated multi-layered immunity.

## Introduction

Prokaryotes evolved a highly diverse array of antiviral defence mechanisms^1–3^. These defence systems function not only as weapons in the evolutionary arms race with viruses but also as mediators of internal genetic conflicts. Within a single microbial cell, mobile genetic elements, such as prophages, phage satellites, conjugative elements, and plasmids, have divergent “interests”, leading to multilateral conflicts that shape cell fates^4^.

Many components of prokaryotic immune systems are evolutionarily linked to eukaryotic innate immunity^5–10^. Multiple key immune genes diversified in prokaryotes and were later vertically inherited by or horizontally transferred to eukaryotes, contributing to the emergence of innate immune pathways^6^. Among the most ancient and versatile components of immune systems in both prokaryotes and eukaryotes are P-loop NTPases^11^. These NTPases comprise one of the most abundant protein folds in all three domains of life as well as in viruses and couple ATP or GTP hydrolysis to an enormous variety of biological processes. Defence systems primarily encompass two types of P-loop NTPases, helicases and signalling NTPases. Most signalling NTPases involved in defence belong to the STAND superfamily (signal transduction ATPases with numerous domains)^12^. Members of the STAND superfamily, such as Apaf1 and CED-4, as well as diverse plant NLR (Nucleotide-binding Leucine-rich Repeats) ATPases, are central components of the programmed cell death machineries in eukaryotes^12^. Although early work proposed that bacterial and archaeal STAND members perform similar roles^13^, for two decades, their functions remained uncharacterised. Recently, however, it has been demonstrated that, despite extensive domain architecture diversity, numerous bacterial STANDs are defence systems that operate via a conserved framework. These NTPases, denoted Avs (antiviral STANDs), are large proteins with a characteristic, tripartite architecture, consisting of a C-terminal sensor domain, middle NTPase domain and N-terminal effector domain. The sensor domain typically forms a repetitive structure, such as an array of a TPRs (tetratricopeptide repeats), which recognises a structural pattern in viral proteins and triggers NTPase-driven oligomerisation and activation of the effector domain^10,14,15^. The effectors of bacterial STANDs, most commonly nucleases, but in some cases, proteases or pore-forming membrane proteins, induce various forms of cell damage that leads to dormancy or cell _death_6,10,14.

KAP (Kidins220/ARMS and PifA) constitutes another family of membrane-bound P-loop ATPases found in both eukaryotes and prokaryotes, characterised by the presence of four transmembrane helices (TMH) located outside of the P-loop domains^16^. Members of this family are involved in signalling and cell defence. A well-characterised representative is the F-plasmid encoded protein PifA, which provides protection against T7^17,18^, but despite sharing the KAP family fold, lacks TMHs^16^. As with STAND systems, KAP ATPases often contain structural repeats and accessory domains that are thought to perform sensing and effector functions^16^.

In this work, we focus on Tmn, a member of the YobI family of P-loop NTPases^19^. A signature feature of this family, absent in other P-loop NTPases, is the insertion of two TMH within the P-loop domain itself^11,16^, which anchors the NTPase domain at the membrane and likely couples ATP hydrolysis to membrane-associated functions^16,20^. In contrast to STAND or KAP, YobI family members lack readily identifiable accessory domains, complicating functional characterisation. Tmn has recently been implicated in anti-phage defence and shown to synergise with ATP-sensitive defence systems such as Gabija, Septu type I, and PrrC^19,21^, but its mechanisms of action and coordination with other defence systems remained unknown.

Here, we show that Tmn is a decameric transmembrane protein that senses phage infection via a solenoid-like cytosolic arm composed of tandem α-helical repeats. These repeats are evident in 3D models despite weak sequence-level signatures and encompass HEAT- and ARM-like motifs, suggesting that Tmn contains a previously unrecognised sensor scaffold involved in bacterial immunity. Phage sensing occurs through direct interaction of this repetitive domain with the phage T2 protein RIIB, which is involved in phage DNA replication^22^. Once activated, the ATPase domain of Tmn drives rapid magnesium ion export, creating a hyperosmotic imbalance that causes plasmolysis (i.e., cytoplasmic shrinkage) and arrests phage development. We further uncover the molecular basis of Tmn synergy with ATP-depletion-sensing defence systems, showing that Tmn-driven ATP hydrolysis serves as a metabolic trigger that activates Gabija and Septu type I, enhancing protection and limiting secondary phage spread. Thus, Tmn functions not only as a self-contained defence system, but also as an upstream trigger that links membrane-localised phage sensing to broader immune activation.

## Results

### Phylogenomics and functional diversification of Tmn

Tmn was identified in multiple prokaryotes^19^, and in *E*. *coli* is often located in F-type plasmids^21^. To investigate the full diversity of Tmn in prokaryotes, we examined precomputed PADLOC results for 223,572 bacterial and archaeal genomes^23,24^. We identified Tmn defence systems in 5,915 genomes across diverse bacterial groups including Pseudomonadota (68% of the occurrences), Bacillota (24%) and Bacteroidota (7%), as well as a single archaeal genome (**Supplementary Table 1**). This distribution suggests that Tmn can operate across a wide range of cellular envelopes and genomic contexts, rather than being restricted to a particular lineage or cell wall or membrane type.

Following MMseqs2 clustering, we compiled a representative set of 785 Tmn sequences to describe system diversity. Multiple sequence alignments confirmed the conservation of signature Walker A and Walker B motifs and two TMHs embedded within the P-loop NTPase domain^16^, whereas the C-terminal region, dominated by predicted α-helices, exhibited considerable variability (**Fig. 1a,b**).

**Figure 1.**
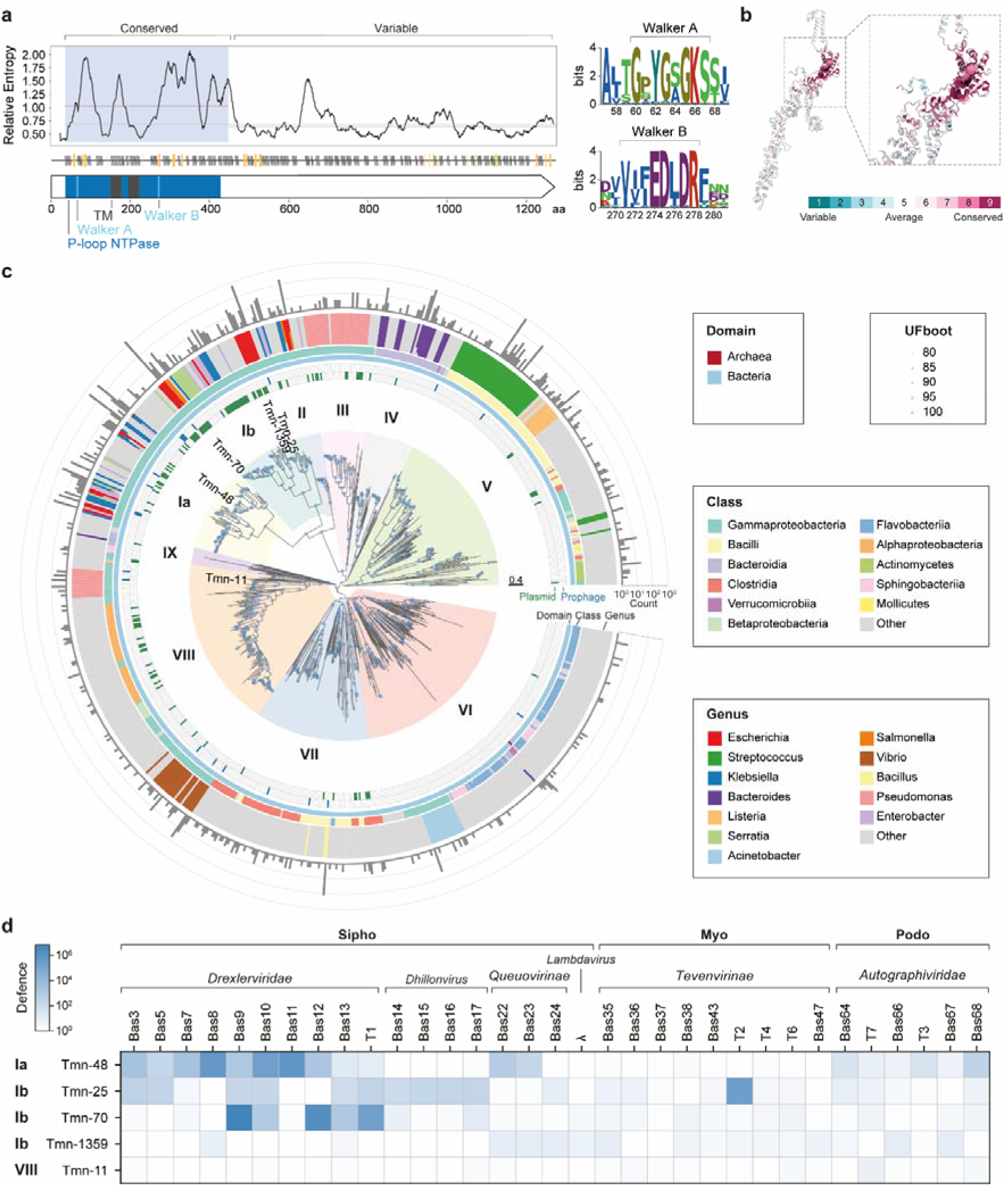
Tmn is phylogenetically diverse and provides protection against diverse phages. a, Conservation and domain architecture of Tmn proteins. Line plot represents relative entropy along the protein sequence, indicating conserved (high relative entropy) and variable (low relative entropy) regions. The transmembrane (TM) domains and P-loop NTPase motifs (Walker A and Walker B) are highlighted. Sequence logos of Walker A and Walker B motifs are shown to the right of the line plot. **b,** Structural conservation of Tmn determined by ConSurf. Tmn protein is coloured by sequence conservation. Inset shows a zoomed-in view of the conserved core region. **c**, Phylogenetic distribution of Tmn proteins. Phylogenetic tree of Tmn homologues, grouped into 10 clades (Ia-IX). The inner rings show Tmn associated with plasmids (green) and prophages (blue). The middle rings show the bacterial domain, class and genus colour-coded according to the key on the right. The outer ring shows the prevalence (count) of each Tmn homolog. Branches with bootstrap values (UFBoot) above 85% are indicated with circles according to the key on the right. **d,** Tmn provides defence against diverse phages. Heatmap shows defence activity of Tmn variants from different bacterial hosts against taxonomically diverse phages. Defence levels are indicated by colour intensity.

Phylogenetic analysis identified 10 major clades (Ia-IX) of Tmn proteins, characterised by distinct protein lengths, genomic contexts, and taxonomic distribution (**Fig. 1c**, **Fig. S1a**). Analysis of the genomic neighbourhoods did not identify a conserved operon architecture, consistent with Tmn functioning as a self-contained defence unit that can appear in diverse genomic contexts (**Supplementary Table 2**). Closely related sister clades Ia and Ib have similar protein lengths (median ∼1270 aa) and are found exclusively in *Enterobacteriaceae* (Gammaproteobacteria), but members of clade Ia are enriched on chromosomes, whereas members of clade Ib are enriched on plasmids (**Fig. 1c**). Other clades show distinct lineage biases and length distributions (**Fig. 1c**, **Fig. S1a**), consistent with repeated diversification of the variable C-terminal region. Notably, 25 genomes encoded two Tmn homologues (**Fig. S1b**). Although co-occurrence alone does not prove distinct specificity, it is consistent with the possibility that multiple Tmn variants broaden defensive range. In most instances, co-occurring Tmn systems belonged to the same major clade and were in two cases located in neighbouring leaves of the phylogenetic tree, indicating that even variants with limited differences can co-exist and might differ in functional output (**Fig. S1b**).

Variability among and within Tmn clades is concentrated in the C-terminal domain, suggesting that this region encodes determinants of the targeting range (**Fig. 1b**). To test this prediction, we selected five representative Tmn variants: one from clade Ia (chromosomally-encoded), three from clade Ib (typically plasmid-encoded), and one from the more distant clade VIII (**Fig. S1c**). These variants were tested for their ability to protect *E. coli* against a panel of 33 diverse phages (**Fig. 1d**). As anticipated, each variant displayed a unique protection profile, with differences observed even among closely related Ib proteins, supporting the possibility that subtle sequence (and inferred structural) variation, particularly in the C-terminal region of Tmn, tunes target recognition (**Fig. 1d**).

Together, these findings show that Tmn is a phylogenetically diverse family of antiviral proteins, with multiple clades differing in genomic context, taxonomic distribution, and phage specificity. The C-terminal variability of Tmn proteins is likely to be the key determinant of the specificity of trigger recognition and thus could play an important role in shaping the defensive repertoire of the system.

### Tmn is a decameric transmembrane complex with a solenoid-like C-terminal domain

To elucidate the oligomeric organisation of Tmn, we combined single-particle cryo-electron microscopy (cryo-EM) with AlphaFold 3 (AF3)-guided modelling. Cryo-EM analysis yielded a three-dimensional reconstruction at an overall resolution of ∼11.6 Å after non-uniform refinement with C5 symmetry was applied (**Fig. S2a,b**). Despite extensive optimisation of sample preparation and cryo-EM processing, including testing alternative expression systems, detergent screening to reduce preferred-orientation bias, stringent exposure curation, and multiple rounds of 2D and 3D classification/heterogeneous refinement, pronounced particle heterogeneity and limited high-resolution features precluded atomic model building (**Fig. S2b**, Methods).

For visualisation, the reconstruction is shown with an AF3-guided model fitted into the density (**Fig. 2a**). The resulting architecture reveals a highly ordered decamer composed of 10 elongated protomers arranged radially around a central axis. This oligomeric state is distinct from the hexameric rings typical of numerous P-loop NTPases^25^, as well as from the tetrameric assemblies described for many Avs^14^. In side view, the Tmn decamer adopts an umbrella-like organisation with an overall vertical height of ∼251 Å, whereas the top view shows a flower-shaped arrangement with an apical diameter of ∼68 Å and a basal diameter of ∼287 Å (**Fig. 2a**). The protomers converge towards a compact central hub, forming a continuous axial channel that traverses the assembly. Measurements from the map indicate that the narrowest constriction of this channel is ∼25 Å in diameter, consistent with a defined central opening (**Fig. 2a**). Given the pronounced heterogeneity observed on grids, we interpret the reconstruction as capturing a predominant oligomeric state that might coexist with additional conformations in solution.

**Figure 2.**
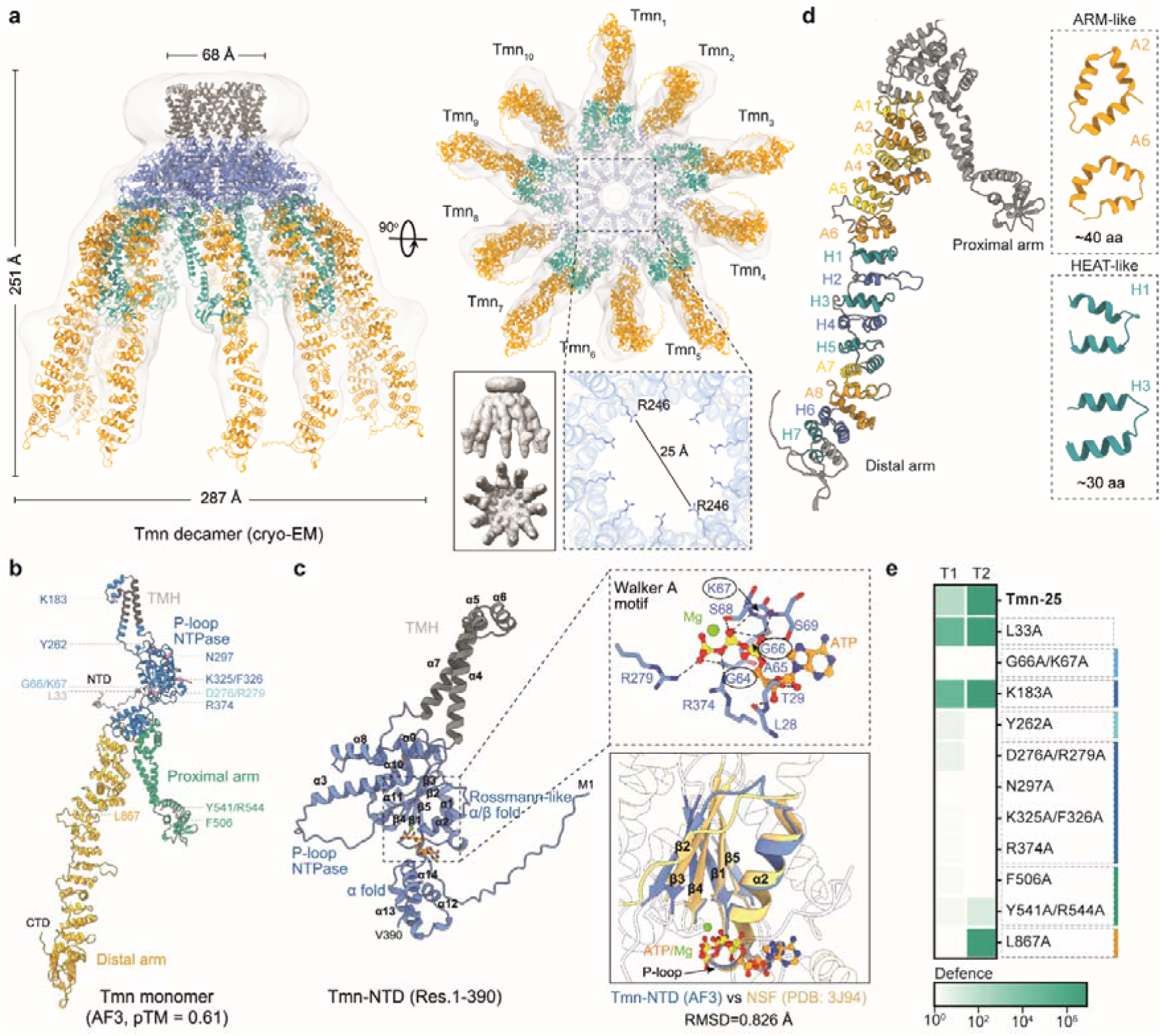
Spatial architecture and repeat organisation of the Tmn-25 defence system. **a**, Cryo-EM reconstruction of the Tmn decamer at ∼11.6 Å resolution, shown as semi-transparent density surface and overlaid with an Alphafold 3 (AF3)-guided-predicted hybrid model (cartoons), shown in side and top views. The decamer forms a prominent central axial channel that is lined, in the fitted model, by inward-facing P-loop NTPase domains from each protomer. A zoomed-in view highlights the ordered arrangement of these domains around the channel, defining a pore at the centre of the assembly. **b**, AF3-predicted Tmn monomer, coloured by structural regions. NTD, N-terminal domain; CTD, C-terminal domain; TMH, transmembrane helices. Amino acid mutations tested in (e) are mapped onto the structural model. **c,** Structural organisation of the Tmn N-terminal region (residues 1-390). The Tmn P-loop NTPase domain adopts a conserved Rossmann-like α/β fold. Top right, inset shows the AF3-predicted Walker A region and a model of Mg^2+^-ATP coordination. Bottom right, structural superposition of the Tmn N-terminal domain with the N-ethylmaleimide-sensitive factor (NSF, PDB: 3J94), a well characterised AAA+ ATPase, reveals a high degree of structural similarity (RMSD = 0.826 Å). **d**, The distal arm of Tmn adopts a modular solenoid-like architecture composed of eight ARM-like repeats (A1-A8, yellow and orange) and seven HEAT-like repeats (H1-H7, blue and teal). Left: repeats mapped onto a single Tmn monomer with cropped P-loop NTPase. Right: example of the ARM-like and HEAT-like repeat structures. **e**, Functional impact of mutations on phage defence, shown as a heatmap of efficiency of plating (EOP) against phage T1 and T2.

To interpret the low-resolution density, we integrated AF3 predictions with the experimental reconstruction. AF3 modelling of the Tmn N-terminal region supported higher-order assembly, whereas full-length multimer predictions were limited to lower-order states (dimers), due to the current constraints in structural prediction for large assemblies (**Fig. S2c,d**). We therefore generated a hybrid full-length decamer model by combining AF3-predicted oligomeric and monomeric models through superposition of the transmembrane helices and adjustment of the short connecting regions, followed by rigid-body refinement against the cryo-EM density (**Fig. 2a**). At the achieved resolution, this approach does not resolve atomic interactions but provides a coherent placement of major domains and the overall assembly geometry.

Each Tmn protomer is anchored in the membrane by the two TMH and contains an N-terminal P-loop NTPase domain, followed by extended cytosolic regions that project outward from the decameric hub (**Fig. 2a**). We refer to these α-helical extensions as proximal and distal arms to reflect their relative positions and lengths (**Fig. 2b**). The N-terminal NTPase domain adopts the typical Rossmann-like α/β-fold, with the Walker A and B motifs juxtaposed in the cytosol, and Walker A forming the ATP-binding pocket (**Fig. 2c, top right**). Structural superposition with the ATPase N-ethylmaleimide-sensitive factor (NSF) reveals close correspondence of the α/β core and Walker A geometry (RMSD = 0.826 Å), indicating a conserved mode of ATP engagement despite the distinct higher-order organisation of Tmn (**Fig. 2c, bottom right**). Whereas NSF couples nucleotide binding and hydrolysis to ring-based mechanochemical cycling within hexameric assemblies^26,27^, Tmn embeds a closely related nucleotide-recognition core within a decameric transmembrane hub, suggesting that conserved ATP-binding and hydrolysis chemistry can be deployed in fundamentally different oligomeric scaffolds.

Within the Tmn decamer, protomers associate as dimers that serve as the fundamental building blocks of the assembly, with the decamer organised as a pentamer of dimers. The intra-dimer interface is extensive and involves coordinated contacts between the transmembrane helices, P-loop NTPase domains, and proximal arm regions, stabilised by complementary electrostatic interactions and hydrophobic packing (**Fig. S2d**). Comparison with AF3-predicted Tmn dimers shows that the experimentally observed dimer closely resembles the predicted architecture, except for the relative positions of the proximal and distal arm regions (**Fig. S2e**). In the cryo-EM model, the proximal arms adopt a more compact configuration relative to the distal arms and approach one another more closely within the dimer, forming additional contacts that are not fully captured in the AF3 dimer prediction. The dimers are connected through inter-dimer interfaces that share some of the intra-dimer interactions but additionally involve surface-exposed regions of the P-loop NTPase domains and adjacent arm segments not engaged in the intra-dimer interface (**Fig. S2f**). Despite these distinct interface contexts, superposition of protomers from intra- and inter-dimer arrangements reveals nearly identical core conformations (RMSD = 0.640 Å), with only modest structural variability confined to α-helical elements of the NTPase domain and the proximal and distal arm regions (**Fig. S2f**, right). Thus, the decamer is built by repeating a largely invariant protomer core around the ring, with only minor flexibility in peripheral helices and arms, rather than by major conformational changes during assembly. Notably, the local differences between AF3 predictions and the cryo-EM-guided model are concentrated in the peripheral arm regions, consistent with intrinsic flexibility and/or context-dependent rearrangements.

Because many P-loop NTPases associate with tandem α-helical repeat domains, including TPR and leucine-rich repeat (LRR) motifs^10,12,14,15^, we searched the proximal and distal arms of Tmn for internal repeats. Repeat analysis tools (REP2^28^, HHrepID^29^ and STRPsearch^30^) did not identify any high-confidence conserved repeats in either region. Nonetheless, AF3 models for all major Tmn clades consistently predict that the distal arm adopts an α-solenoid fold comprising two repeat-like elements: the upper and lower segments contain Armadillo (ARM)-like repeats, each formed by three α-helices, whereas the central segment contains Huntingtin, elongation factor 3, protein phosphatase 2A, TOR1 (HEAT)-like repeats consisting of two antiparallel helices (**Fig. 2d** and **Fig. S2g**). The overall architecture and repeat number are highly conserved across representatives of all major Tmn clades (Icarus^31^ TM score 0.997 – 0.807), with six ARM-like repeats in the upper segment, five HEAT-like repeats in the central region, and two ARM-like and two HEAT-like repeats in the lower segment, although individual proteins vary in loop length and in the size of the proximal α/β C-terminal domain (**Fig. S2g**). We note that some homologues (e.g. WP_152187733.1) with longer distal arms are predicted to contain additional ARM-like elements, which correlate with extended proximal arm regions, consistent with modular expansion of the arm architecture.

This arrangement is consistent with a solenoid architecture, in which repetitive α-helical units stack into a curved or superhelical array^32^. α-solenoids often function as scaffolds for protein-protein interactions and are common in regulatory and transport proteins^32,33^. In Tmn, the juxtaposition of the more rigid ARM-like elements with the more flexible HEAT-like elements suggests a structural basis for bending within the central region, potentially providing a tunable scaffold for engaging the recognised structures. By contrast, the proximal arm is predicted to form a conserved coiled-coil attached to a variable α/β domain that differs in length and shape across Tmn homologues (**Fig. S2g**).

To functionally probe the structural model, we performed targeted mutagenesis of predicted functional residues in Tmn-25 (**Fig. 2e**). Mutations in the conserved Walker A motif (G66A/K67A) abolished defence activity. Additional substitutions in the YobI-like P-loop NTPase domain (Y262A, D276A/R279A, N297A, K325A/F326A, and R374A) also disrupted function (**Fig. 2e**). Based on the model, these residues cluster around the predicted nucleotide-binding pocket and adjacent structural elements, consistent with roles in Mg^2+^ coordination (D276, R279), stabilising nucleotide binding (K325), or coupling nucleotide state to conformational change (Y262, N297, F326, R374). These mutational phenotypes indicate an essential requirement for nucleotide binding and hydrolysis for Tmn activity.

Mutations in conserved residues of the proximal arm (F506A, Y541A/R544A) also abolished defence, consistent with a critical role of this region in structural integrity and/or protein-protein interaction (**Fig. 2e**). A mutation in the distal α-solenoid domain (L867A) located within the ARM-like repeat A5 selectively abolished defence against phage T1 while retaining protection against T2, suggesting that distinct elements within the distal arm can influence the antiphage specificity of Tmn (**Fig. 2b,d,e**). By contrast, mutations of poorly conserved residues in the N-terminal region (L33A) and in the P-loop NTPase domain (K183A) had no negative impact on Tmn activity (**Fig. 2e**).

In summary, Tmn forms a membrane-anchored, decameric defence complex comprising conserved NTPase cores organised in a central hub and extended cytosolic α-helical domains with solenoid-like features. Sequence and structural variation within the proximal and distal arms across Tmn clades, together with the distinct phenotype of a distal-arm mutant, supports a model in which these regions act as sensors that bind target proteins and determine defence specificity.

### Tmn disrupts phage replication by inducing plasmolysis of the infected cell

Membrane-associated P-loop NTPases have been proposed to regulate NTP-dependent assembly or disassembly of membrane-associated protein complexes^16^. A relatively well-characterised example is PifA, a KAP family member that defends bacteria against phage T7 by triggering membrane depolarisation and nucleotide leakage, resulting in abortive infection^17,18,34^. Although Tmn belongs to the phylogenetically distinct YobI family of P-loop NTPases, it similarly anchors to the membrane, so we hypothesised that it could employ a similar defence mechanism.

Structural analysis revealed that the Tmn decamer forms a central pore of approximately 25 Å (2.5 nm), large enough to permit the passage of small molecules such as nucleotides (ATP ∼1 nm). To test whether Tmn mediates nucleotide leakage during infection, we measured total and extracellular ATP levels in Tmn-expressing cells and YFP-expressing controls infected with phage T2. While the total ATP level dropped substantially in Tmn-expressing cells post-infection, extracellular ATP level remained unchanged (**Fig. 3a**), indicating that ATP is being hydrolysed intracellularly rather than leaking. This finding suggests that, in response to phage infection, the P-loop NTPase of Tmn exhibits strong ATPase activity, sufficient to cause a measurable drop of the ATP level, rather than acting as a nucleotide channel. This was further confirmed by fluorescence-based permeability assays to assess membrane integrity. Using a fluorescence plate reader, we monitored propidium iodide (PI) uptake, indicative of membrane damage, and DISC(3)5 fluorescence, a marker for membrane depolarisation. Unlike PifA, Tmn-expressing cells showed no signs of membrane depolarisation or increased permeability at any multiplicity of infection (**Fig. S3a**). These findings were further validated by fluorescence microscopy: in control cells, phage T2 infection led to PI entry, consistent with phage-induced membrane damage. In contrast, infected Tmn-expressing cells resisted PI entry, and FM4-64 staining showed an intact membrane (**Fig. 3b,c**).

**Figure 3.**
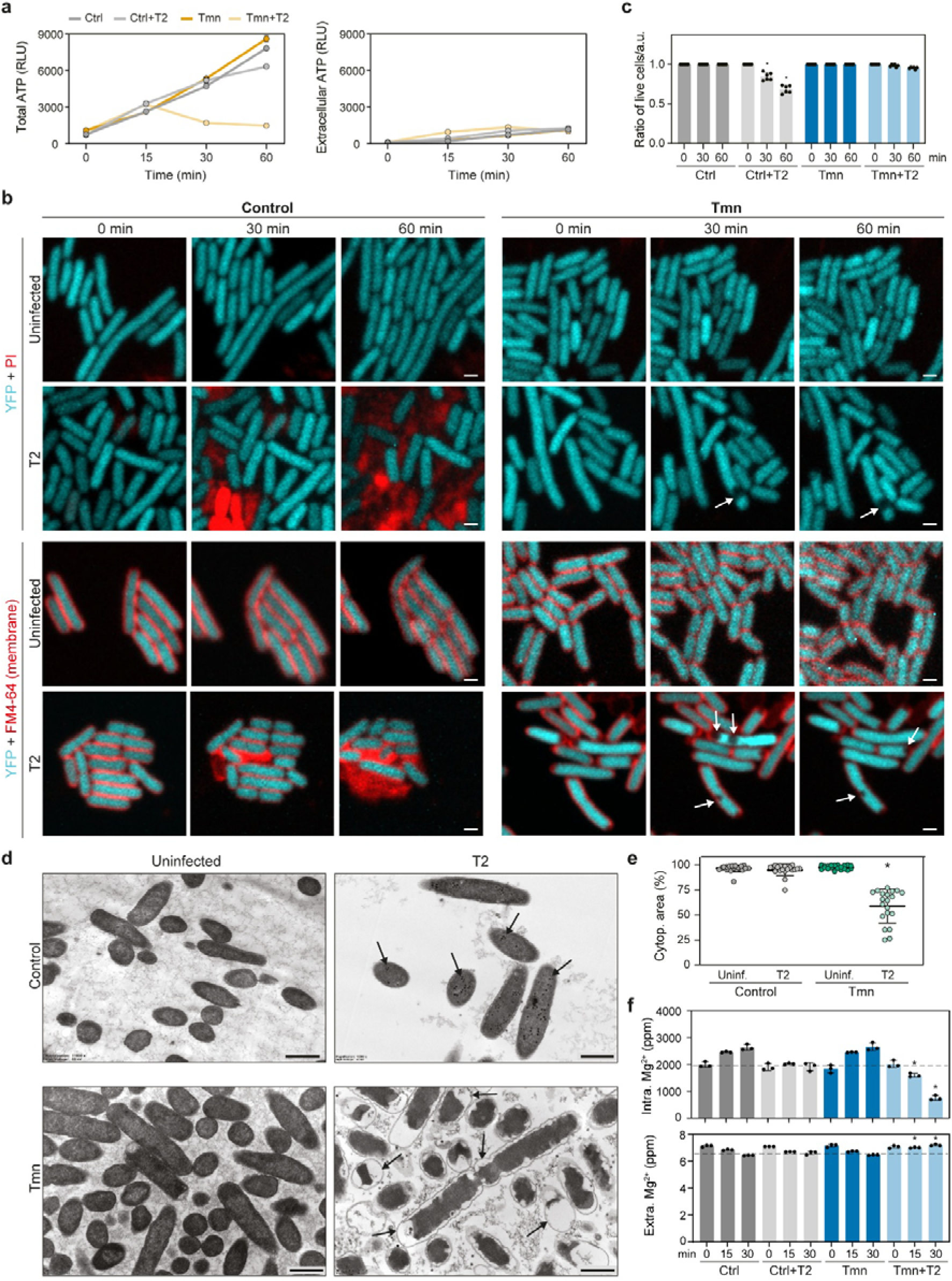
Tmn provides anti-phage defence via plasmolysis. **a**, Quantification of total (left) and extracellular (right) ATP levels over time in control (YFP-expressing) and Tmn-expressing cells, infected or uninfected with phage T2 (MOI ≈ 10). **b**, Time-lapse confocal microscopy assessment of viability of control (YFP) and Tmn cells uninfected and infected with phage T2. Viability was assessed with propidium iodide (PI) (top) and membrane was stained with FM4-64 (bottom) over 60 min. Arrows indicate cytoplasmic holes observed in infected Tmn cells. Scale bar, 1 µm. **c,** Quantification of live cells of uninfected and infected control (YFP) and Tmn cells, visualised by fluorescence microscopy. Asterisks indicate significant differences (p < 0.05) in comparison to the initial time point of the same condition. **d**, TEM images of cross-sections from uninfected and infected control (YFP) and Tmn cells. In infected control cells, arrows point towards phage capsids formed inside the cells. In infected Tmn cells, arrows point towards examples of detachment of inner membrane from the outer membrane (plasmolysis). **e**, Quantification of cytoplasmic area as a percentage of total cell area from TEM cross-sections. The relative cytoplasmic area of 20 cells over three independent images are depicted for each condition. Asterisk indicates significant differences (p < 0.05) of Tmn cells in comparison to all other conditions. **f**, Quantification of intracellular (left) and extracellular (right) magnesium (Mg^2+^) levels measured over time by electrospray ionisation mass spectrometry in uninfected and infected control and Tmn cells. Asterisks indicate significant differences (p < 0.05) of Tmn cells in comparison to the respective control cells.

However, microscopy revealed a surprising phenotype in infected Tmn-expressing cells, with the appearance of discrete cytoplasmic “holes”, regions of the cell devoid of YFP signal (**Fig. 3b**, white arrows). To explore these regions, we used cross-sectional transmission electron microscopy (TEM) of infected cells. In controls, mature T2 capsids were visible by 15 min post-infection, indicating normal phage development. In contrast, Tmn-expressing cells lacked mature capsids and showed signs of cytoplasmic shrinkage and detachment of the inner membrane from the outer membrane, hallmarks of plasmolysis^35^ (**Fig. 3d**). Quantification of the cytoplasmic area relative to the total cell area confirmed a ∼40% reduction in infected Tmn cells compared to all other conditions (**Fig. 3e**).

Plasmolysis typically occurs in response to hyperosmotic stress, where water efflux via osmosis leads to cytoplasmic shrinkage and membrane detachment^35^. Given that we observed high level of ATP hydrolysis by the ATPase domain of Tmn upon phage infection but no membrane leakage, we hypothesised that Tmn induces plasmolysis by actively exporting ions to create osmotic imbalance. To test this hypothesis, we measured intra- and extracellular concentrations of potassium (K^+^), calcium (Ca^2+^), sodium (Na^+^), and magnesium (Mg^2+^) at 0-, 15-, and 30-minutes post-infection. The Mg^2+^ level showed a marked Tmn-dependent change, with infected Tmn-expressing cells exhibiting substantially decreased intracellular and increased extracellular Mg^2+^, consistent with Mg^2+^ export upon Tmn activation (**Fig. 3f**, **Fig. S3b**). Changes in the Na^+^ and K^+^ levels were also observed during infection but did not depend on Tmn activation (**Fig. S3b**), indicating that the perturbations of these ions reflected broader, infection-associated homeostatic responses. These findings indicate that Tmn protects infected cells by coupling ATP hydrolysis to selective Mg^2+^ export, thereby inducing plasmolysis.

### Tmn-induced plasmolysis delays lysis and blocks phage transmission

To assess how Tmn-induced plasmolysis affects infection dynamics, we used a microfluidic platform adapted for monitoring single-cell infections^36^ (**Fig. 4a**). This system allows controlled infection at a MOI ≈ 1 and real-time tracking of cell growth, lysis, and lineage expansion.

**Figure 4.**
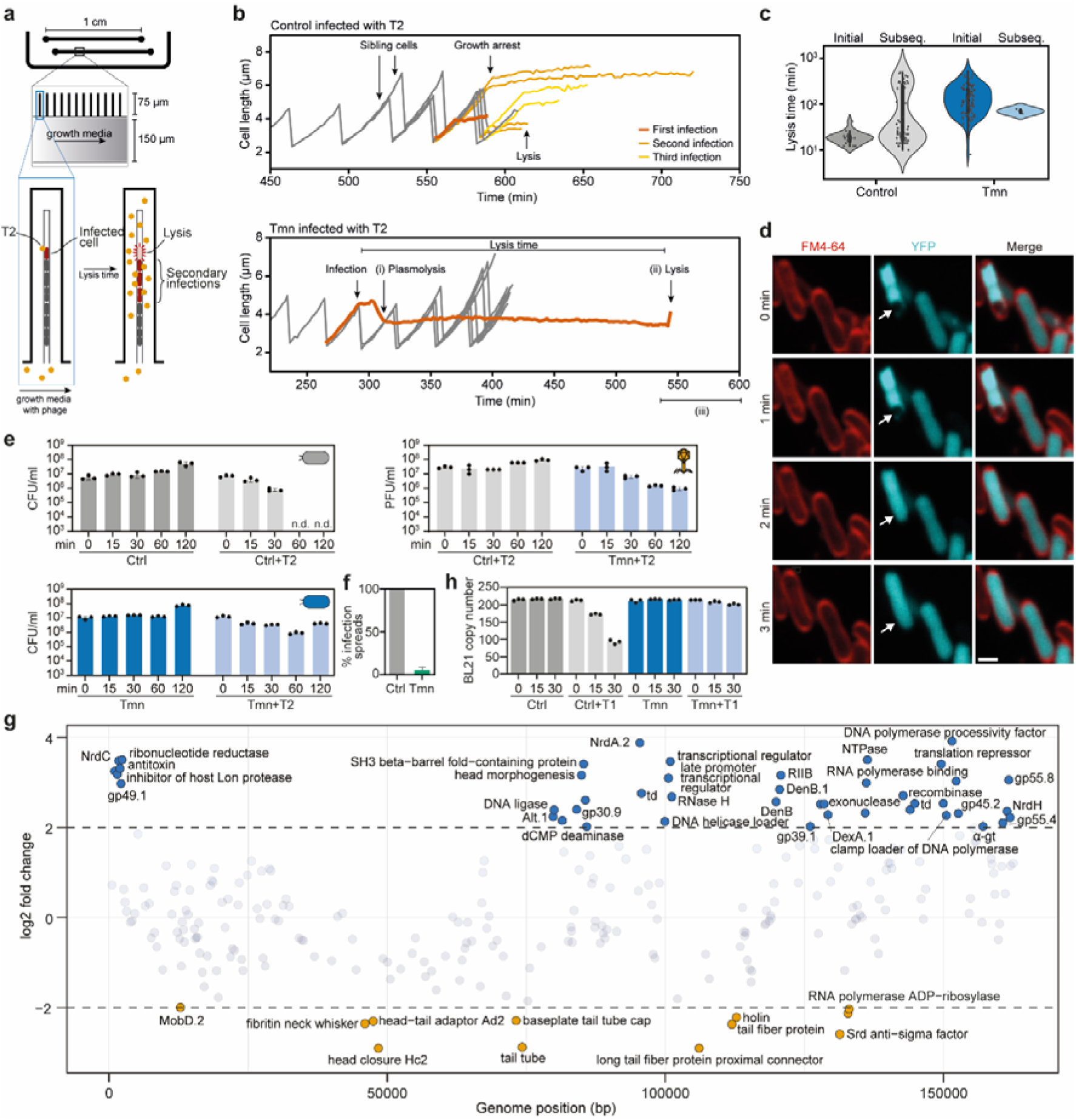
Tmn-induced plasmolysis is reversible and blocks phage propagation before the cell is irreversibly damaged. **a**, Schematic of the microfluidic platform used for real-time, single-cell analysis of phage infection. Bacteria are confined in trenches where they grow clonally, allowing precise delivery of phage and continuous imaging. **b,** Control cells (top) exhibit abrupt growth arrest and lysis following infection, followed by infection spread and lysis inhibition events. Tmn cells (bottom) undergo (i) plasmolysis shortly after infection, followed by (ii) delayed lysis while sibling cells proliferate (iii). **c,** Violin plot of lysis times for first and subsequent infection events in control versus Tmn-expressing cells (n = 165 total infection events). **d,** Time lapse confocal microscopy of Tmn-expressing cells during phage T2 infection, stained with FM4-64 (membrane) and expressing cytoplasmic YFP. Arrows highlight cytoplasmic holes disappearing over time. Time is represented in relation to the appearance of a cytoplasmic hole, indicated as time 0. Scale bar, 1 µm. **e,** Time post infection assays measuring bacterial (CFU/ml) and phage (PFU/ml) concentrations for T2 infected and uninfected control (YFP) and Tmn cells. n.d., not detectable. **f**, Quantification of infection spreads to neighbouring cells during T2 infection of control and Tmn-expressing cells, measured by single-cell microscopy. **g**, RNAseq analysis of phage gene transcription levels in cells harbouring Tmn in relation to control cells, represented as log2 fold differences. Phage genes with upregulated transcription in Tmn cells are coloured blue, while those downregulated are coloured orange. **h**, Host chromosome abundance during infection of control and Tmn-expressing cells with phage T1. Bars show the estimated mean BL21 genome copy number (derived from genome-wide sequencing read coverage, used here as a proxy for host DNA abundance) and dots indicate independent biological replicates.

In control cells, T2 infection caused rapid growth arrest followed by cell lysis approximately 20 min later (**Fig. S4a**). Released phages then spread to neighbouring cells within ∼6 min, as indicated by the immediate growth arrest of adjacent cells (**Fig. 4b, Fig. S4a**). These secondary infections often occurred at high MOIs, triggering lysis inhibition^36^ in some cells, manifested as substantially delayed lysis (mean ∼300 min), resulting in a bimodal distribution of lysis times (**Fig. 4c**).

In contrast, Tmn-expressing cells consistently underwent plasmolysis shortly after infection, manifested as a sharp reduction in cell length (**Fig. 4b**, blue arrows in **Fig. S4a**). Lysis was delayed and phage progeny was not released, preventing further infections within the trench (**Fig. 4a,b, Fig. S4a**). Across 165 infection events (76 control, 86 Tmn), Tmn expression consistently delayed lysis, allowing sibling cells to grow and divide (**Fig. 4b**). Thus, Tmn effectively uncouples infection from phage transmission, halting phage replication without arresting population growth (**Fig. S4b**).

In summary, Tmn enables infected cells to isolate phage infection through plasmolysis, allowing the uninfected siblings to survive and expand, thereby protecting the bacterial population.

### Tmn stalls progression of infection at a reversible stage

Given that plasmolysis can be a reversible stress response^35^, we asked whether Tmn-expressing cells could recover once the phage threat was neutralised. Fluorescence microscopy revealed that the cytoplasmic “holes” characteristic of plasmolysis gradually disappeared over time, suggesting that Tmn-induced cytoplasmic collapse can resolve rather than representing an obligate terminal state (**Fig. 4d**). Bulk infections at high MOI showed that phage titres decreased due to Tmn-mediated defence, and colony-forming units (CFUs) declined less in Tmn-expressing cells compared to controls, with a measurable surviving fraction (7%; 9x10^5^ out of 1.3x10^7^ CFUs/ml) (**Fig. 4e**). Thus, a subset of cells can outlast the infection-associated stress and resume growth once phage propagation is curtailed.

At the single-cell level, the Tmn response sharply limited transmission of infection, with only 4.8% of Tmn trenches showing any secondary infection, and even in these rare cases, only one neighbouring cell was affected, reflecting an extremely low burst size (**Fig. 4f**). This reduction in progeny production was confirmed by bulk one-step growth curves, where T2 achieved a burst of 90±15 PFU/cell in control cells and only 3±1 PFU/cell in Tmn-expressing cells (**Fig. S4c**).

To define the step at which Tmn-induced plasmolysis disrupts the phage infection cycle, we performed RNA-seq on infected cells. Tmn-expressing cells showed a marked reduction in transcription of late phage genes encoding structural components (e.g., neck, baseplate, tail fibre connectors) and lysis factors (e.g., T holin), consistent with the absence of mature capsids observed in cross-sectional TEM (**Fig. 4g**, **Fig. 3d**). Conversely, we detected increased relative abundance of transcripts corresponding to early-to-middle infection functions, including DNA replication and recombination (e.g., RecA, DNA polymerase clamp loader subunit A, 59 protein, gp45.2), nucleotide metabolism (e.g., thymidylate synthase, NrdH glutaredoxin), and multiple regulatory factors for late transcription (e.g., RegA, sliding clamp, PobA, gp55.4, late promoter transcription accessory protein) as well as chaperones such as GroES (**Fig. 4g**). This pattern is most parsimoniously explained by infection-stage arrest, that is, blocking of the phage progression into the late programme, resulting in early-to-middle transcripts remaining abundant because infection does not transition into the late transcriptional state.

Notably, we also detected phage transcripts encoding host DNA degradation factors (e.g., *endoIV/denB* and *dexA*)^37^ (**Fig. 4g**). To test whether Tmn prevents host chromosome degradation during T2 infection, we performed sequencing-based DNA integrity assays. However, under our conditions, T2 phage did not cause detectable host genome degradation even in control cells (**Fig. S4d**). We therefore leveraged phage T1, which robustly degrades the host chromosome in control infections but not in Tmn-expressing cells, indicating that Tmn protection occurs before extensive genome degradation by the phage (**Fig. 4h**). Consistent with this early block, a small fraction of Tmn-expressing cells survived high MOI T1 infection (5% at 30 min; 6.7x10^6^ out of 1.3x10^8^ CFU/ml), comparable to the surviving fraction observed for T2 (**Fig. S4e, Fig. 4e**). Together with the strong suppression of late gene transcription in T2, these data place the Tmn-imposed block at a middle-stage infection checkpoint, after partial phage takeover of the infected cell but before irreversible damage occurs, leaving a window in which some cells can recover. These findings are consistent with a model in which plasmolysis drives rapid physiological arrest that renders the intracellular environment non-permissive for efficient progression of infection.

Together, these results show that Tmn-induced plasmolysis acts as a protective checkpoint that interrupts phage progression. In a minority of the infected cells, this leads to recovery and resumption of growth whereas the majority eventually lyse, but too late for the phage to propagate effectively in the bacterial population.

### Phage protein RIIB directly triggers Tmn activation

To elucidate how Tmn is activated during infection, we isolated and sequenced escape mutants of phage T2 that evaded Tmn-25-mediated defence. All identified escape mutations mapped to *gp317*, which encodes the phage protein RIIB (**Fig. 5a**, **Supplementary Table 3**). RIIB is a phage-encoded replisome component^22^ that is expressed in the early-to-middle stage of infection, prior to DNA replication^38^. The RIIB protein anchors the phage replisome to the host membrane and has also been implicated in membrane remodelling during infection^22^. The convergence of all independent escape mutations in RIIB suggested that it functions as a direct trigger for Tmn activation.

**Figure 5.**
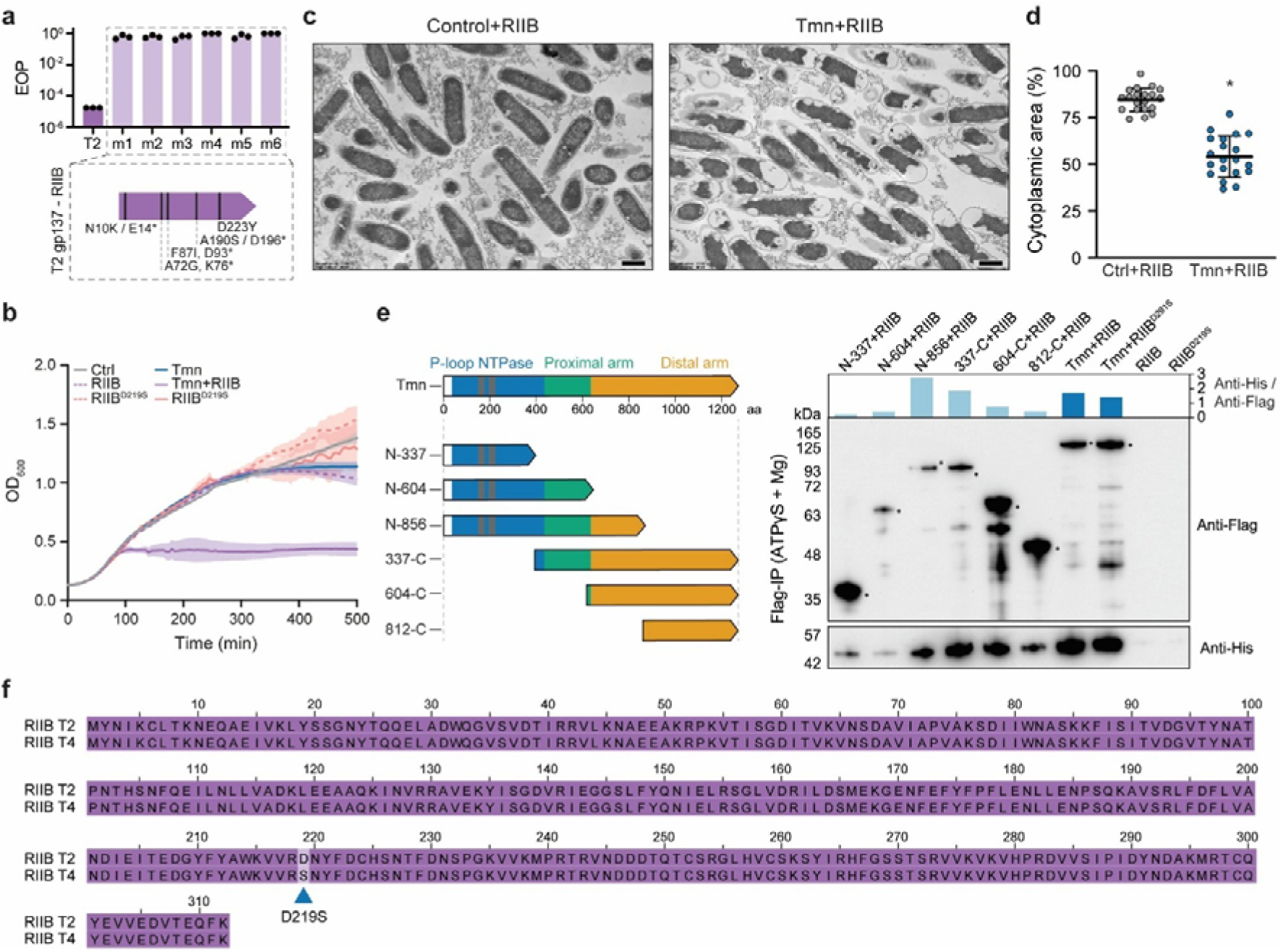
RIIB directly activates Tmn. **a**, Efficiency of plating (EOP) of wild-type phage T2 and six independently isolated T2 escape mutants (m1-m6) on Tmn-25-expressing cells. Lower panel: mutations identified in T2 gp317 (RIIB) across the escape mutants (asterisks denote nonsense mutations). **b**, Growth curves (OD_600_) for control cells, RIIB-only, RIIB^D219S^-only, Tmn-only, Tmn+RIIB co-expression, and Tmn+RIIB^D219S^ co-expression. Lines show mean with shading representing standard deviation. **c,** Representative cross-sectional TEM images of cells expressing RIIB in trans in the absence of phage infection, in a control (YFP) background (left) or in cells expressing Tmn-25 (right). Scale bars are indicated. **d,** Quantification of cytoplasmic area from TEM cross-sections, expressed as a percentage of total cell area. The relative cytoplasmic area of 20 cells over three independent images are depicted for each condition. Horizontal line and whiskers indicate mean ± standard deviation. Asterisk denotes statistical significance (p < 0.05) between Tmn and control cells. **e,** Mapping Tmn regions contributing to recoverable RIIB association by co-immunoprecipitation (co-IP). Left, schematic of full-length Tmn and truncation constructs used. Righ, Flag-tagged Tmn constructs were co-expressed with His-tagged RIIB and subjected to Flag pull-down in the presence of ATPγS and Mg^2+^. Immunoblots were probed with anti-Flag (bait) and anti-His (prey). Bars above the blot report densitometry as the anti-His/anti-Flag signal ratio in the IP for each condition. **f**, Amino acid sequence alignment of RIIB from phages T2 and T4 highlighting a single amino acid substitution (D219S). Purple shading indicates sequenced identity.

To test this prediction, we expressed RIIB *in trans* in cells harbouring Tmn-25. RIIB expression was not toxic to control cells but caused marked toxicity in Tmn-expressing cells (**Fig. 5b**). Cross-sectional TEM further showed that RIIB expression alone was sufficient to induce plasmolysis in Tmn-expressing cells in the absence of phage (**Fig. 5c,d**), demonstrating that RIIB is a direct activator of Tmn-25 rather than a passive marker of the infection stage.

We next asked whether RIIB physically interacted with Tmn and, if so, which portions of the Tmn protein contributed to this association. In co-immunoprecipitation experiments, His-tagged RIIB was reproducibly recovered in Flag-tagged Tmn pull-downs (**Fig. 5e, Fig. S5a**), supporting specific binding. Because RIIB recovery varied across Tmn truncations, we quantified RIIB enrichment by densitometry and normalised anti-His signal to anti-Flag signal in the IP (**Fig. 5e**). This bait-normalised analysis revealed a clear domain dependence. The N-terminal fragment Tmn N-337 (most of the P-loop NTPase) was robustly recovered yet showed near-background RIIB co-precipitation, indicating that the membrane-embedded/NTPase-containing core alone is insufficient for strong association with RIIB. Extending this fragment to Tmn N-604 (adding the proximal arm) did not restore high RIIB recovery, suggesting that the proximal arm alone was not sufficient for high-avidity engagement. By contrast, Tmn fragments that included substantial portions of the cytosolic arms on both sides of the proximal-distal junction (Tmn N-856 and 337-C) showed high anti-His/anti-Flag ratios comparable to those observed with full-length Tmn, indicating that the dominant RIIB-binding determinants reside in the cytosolic arms and are sufficient to recapitulate the Tmn-RIIB association in the absence of the membrane anchor and NTPase core. Distal arm-only fragments (604-C and 812-C) showed detectable but weak RIIB recovery, consistent with a composite, multivalent interface that requires a broader surface than the distal tip alone. Together, these findings indicate that the RIIB engagement by Tmn is multivalent and depends on cooperative contributions from both the proximal and the distal arm regions, but not the NTPase core.

Consistent with these findings, co-expression followed by size-exclusion chromatography yielded partial co-elution of Tmn and RIIB alongside free RIIB and higher-order/aggregated species (**Fig. S5b**), indicating that, under the purification conditions tested, the sample comprises a mixture of complexed and uncomplexed protein populations rather than a unique, homogeneous complex. Similar behaviour was observed in the presence of ATP/Mg^2+^ (**Fig. S5c**), and mass photometry and blue native PAGE likewise revealed heterogeneous species (**Fig. S5d,e**). Thus, our results indicate that RIIB interacts with Tmn but that the complex is not recovered as a single dominant species under the purification conditions tested.

Tmn-25 did not defend against phage T4 which encodes RIIB closely related to the T2 ortholog **(Fig. 1d**). Comparison of T2 and T4 RIIB proteins revealed a single amino acid difference at position 219: an aspartate (D219) in T2 is replaced by a serine (S219) in T4 (**Fig. 5f**). To determine whether this residue was implicated in Tmn activation, we performed co-expression/pull-down assays, where RIIB-D219S remained detectably associated with Tmn, indicating that residue 219 is not strictly required for binding (**Fig. 5e**). Instead, these genetic and functional observations suggested that residue D219 contributes to the efficient activation of Tmn by RIIB, potentially by stabilising a specific interaction geometry or contact required to trigger Tmn response. Consistent with this possibility, growth measurements upon co-expression revealed that RIIB drives a rapid and sustained growth arrest in Tmn-expressing cells, whereas RIIB-D219S does not (**Fig. 5b**). Thus, D219 is dispensable for RIIB binding by Tmn but contributes to sustained activation of the defensive state.

Because the Tmn-RIIB complexes were heterogeneous, we used AF3 modelling to explore plausible RIIB docking configurations on the Tmn assembly. Attempts to model Tmn-RIIB complexes did not converge on a single high-confidence binding configuration, consistent with the transient and context-dependent nature of phage-host activation interactions and with the likely importance of a shared membrane context (both Tmn and RIIB are membrane-associated^39^), which is not explicitly represented in current AlphaFold-based co-folding approaches^40,41^. Nevertheless, across independent modelling attempts, RIIB was repeatedly placed on peripheral arm surfaces, frequently spanning distal and proximal arm-adjacent regions (**Fig. S5f**). This localisation mirrors the co-IP mapping, with the arm-rich constructs that maximise bait-normalised RIIB recovery corresponding to the same peripheral regions most often contacted in AF3 models, providing convergent support for a composite arm interface. This arm-centred interaction model is further supported by comparative sequence analysis of genomes carrying two Tmn proteins. To minimise broad clade-level divergence and focus on substitutions more likely to reflect specificity differences, we restricted this analysis to pairs that occupy neighbouring branches in the phylogeny (that is, closely related variants). In these pairs of closely related Tmn proteins, sequence variation concentrated in the proximal arm (∼30% of the residues substituted, particularly towards its distal end) and distal arm (∼25% of the residues substituted) (**Fig. S5g,h**), supporting a model in which diversification of arm surfaces tunes trigger recognition while preserving a conserved catalytic core.

The RIIB gene is transcribed early in infection and translated during the middle stage, whereas phage enzymes involved in host DNA-degradation, such as EndoII and EndoIV, whose activity results in irreversible cell damage, are transcribed in the middle stage and translated later^38^. This temporal separation suggests that RIIB-dependent Tmn activation occurs before irreversible host damage. This sequence of events is consistent with our transcriptomic data showing that Tmn arrests phage progression at a middle stage (**Fig. 4g**).

In summary, these results identify RIIB as a direct activator of Tmn-25. RIIB expression is sufficient to trigger plasmolysis in the absence of infection, and biochemical assays support specific association between RIIB and Tmn. The requirement for the RIIB residue 219, together with the distinct behaviours of RIIB and RIIB-D219S during co-expression, supports a model in which RIIB activates Tmn through a productive engagement step that is separable from binding and is required to sustain the defensive state.

### The ATPase activity of Tmn triggers synergy with defence systems that sense ATP-depletion

We previously showed that Tmn synergises with several other defence systems, including Gabija, Septu type I, and PrrC^21^. A shared mechanistic feature of these systems is the presence of ATP-sensitive components (GajA, PtuA, and PrrC, respectively) that are inactive or restrained in ATP-replete cells but are activated as the ATP level drops during phage infection^42–45^. Because Tmn displays strong infection-induced ATPase activity (**Fig. 3a**), we reasoned that Tmn could function as an upstream driver of ATP collapse, translating phage infection into an early metabolic signal that de-represses downstream ATP-sensing defences and thereby enables their effector responses.

We first selected phage SLUR435 as a sensitised background for testing synergy because Tmn alone provides weak protection, whereas Gabija and Septu type I alone provide no detectable protection against this phage, but co-expression of Tmn with either Gabija (TG) or Septu type I (TS) resulted in strong protection (**Fig. 6a**). To determine whether Tmn generates the ATP depletion needed for activation of Gabija and Septu, we quantified ATP levels during SLUR435 infection in strains expressing each system alone or in combination. SLUR435 did not cause detectable ATP depletion on its own in control, Gabija or Septu cells, whereas ATP depletion was pronounced in Tmn-expressing cells and remained the dominant signal in the combination strains (**Fig. 6b**). Thus, under these conditions, the phage alone does not generate an ATP signal sufficient to engage Gabija or Septu, whereas Tmn does. Consistent with this conclusion, synergy was abolished in a Walker A mutant of Tmn (G66A) (**Fig. 6a**), indicating that nucleotide binding/hydrolysis by Tmn is required for the synergistic interaction and supporting a hierarchical model of defence activation in which Tmn-driven ATP depletion is the primary upstream trigger.

**Figure 6.**
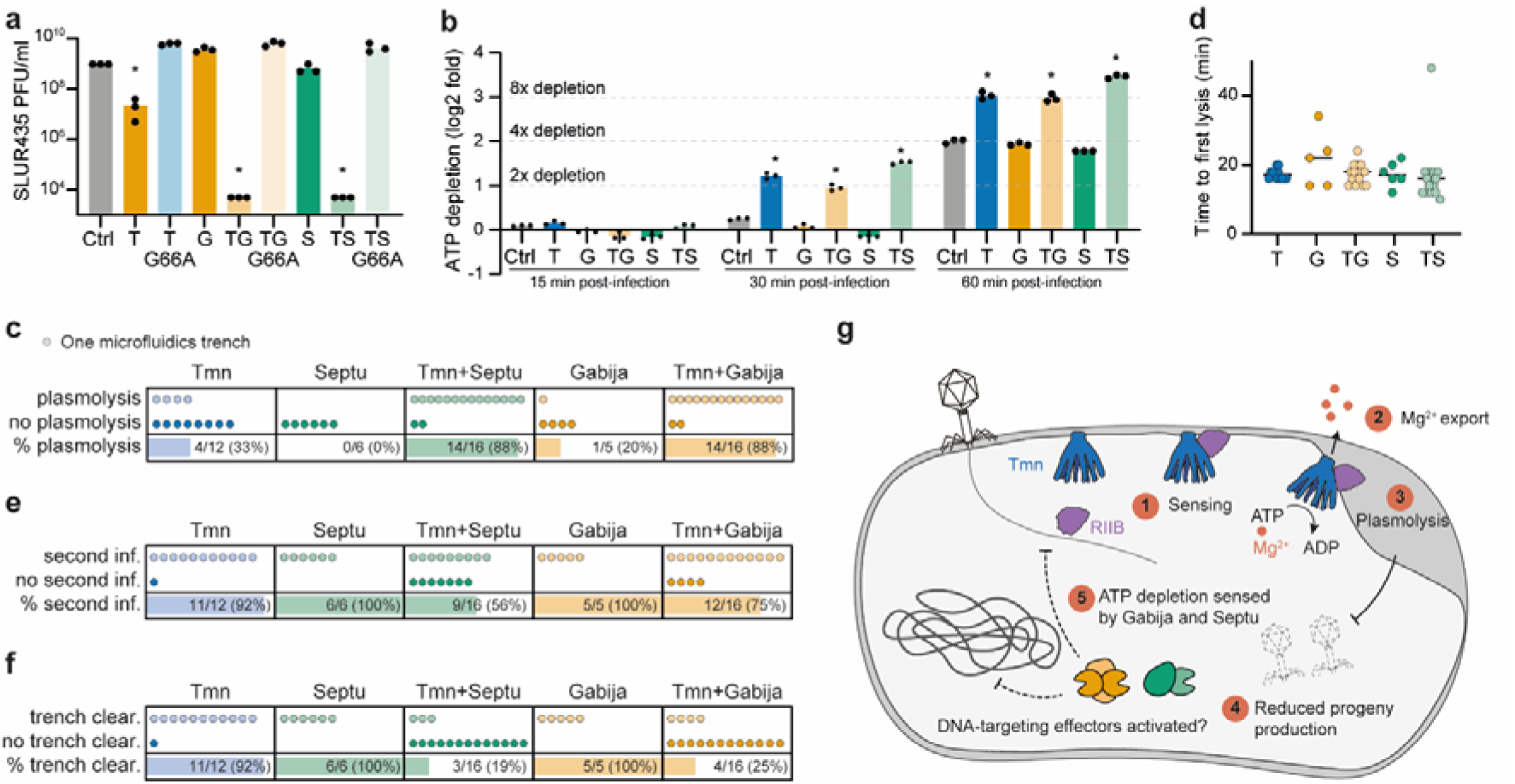
Tmn-driven ATP depletion activates ATP-sensing defence pathways and reshapes infection outcomes. **a**, Synergistic activity of Tmn with Gabija (TG) and with Septu type I (TS) against phage SLUR435, measured by endpoint phage titre (PFU/ml). Walker A mutant controls (G66A) are included to test the requirement for Tmn ATPase activity. Ctrl, control with YFP-expressing cells; T, Tmn; S, Septu type I; G, Gabija. Dots indicate biological replicates and bars indicate mean. Asterisks denote significant differences (p < 0.05) to control. **b,** ATP depletion during infection, plotted as ATP depletion (- Δlog2 fold) in infected cells relative to matched uninfected controls, with each condition normalised to its 0 min value. Higher values indicate greater ATP depletion. Dashed lines indicate 2x, 4x, and 8x depletion. Asterisks denote significant differences (p < 0.05) versus control at the corresponding time point. **c,** Plasmolysis outcomes in microfluidic trenches. Each dot represents one trench scored for whether a plasmolysis event was observed within the imaging window (top row) or not observed (middle row). Bottom bars show the fraction of trenches with plasmolysis (x/N, %). **d,** Time to first lysis in microfluidic trenches. Each point represents one trench; horizontal lines indicate the median time to first lysis. **e,** Secondary infection outcomes in microfluidic trenches. Each dot represents one trench scored for whether a secondary infection event was observed within the imaging window (top row) or not observed (middle row). Bottom bars show the fraction of trenches exhibiting secondary infection. **f,** Trench clearance outcomes following infection. Each dot represents one trench scored for whether immediate trench clearance was observed (top row) or not observed (middle row). Bottom bars show the fraction of trenches with clearance. **g,** Working model for Tmn mechanism of action and ATP-depletion-driven synergy. Left: Phage protein RIIB engages the cytosolic arms of membrane-anchored Tmn, promoting ATP turnover and Mg^2+^ export, leading to plasmolysis and reduced phage progeny production. Right: The ATP collapse generated by Tmn is sensed by Gabija and Septu type I, releasing inhibition of their ATP-dependent sensor modules and enabling activation of their DNA-targeting effectors.

To determine whether this ATP depletion signal propagates into downstream DNA-associated outputs, we used whole-genome sequencing as a population-level readout of host and phage DNA abundance during SLUR435 infection (**Fig. S6a**). At this resolution, we did not detect additional bulk loss of host or phage DNA in Tmn+Gabija or Tmn+Septu relative to Tmn alone. Because sequencing is well suited to detecting fragmentation and copy-number loss but may miss nicking-dominated DNA damage, we assayed DNA integrity by native and alkaline agarose gel electrophoresis (**Fig. S6b**). Under alkaline conditions, nicked DNA is expected to denature and migrate as lower-molecular-weight smears, in contrast to the intact genomic DNA. However, these assays did not reveal a reproducible differential pattern that could unambiguously distinguish additional nicking activity in the combination strains from Tmn alone. Thus, although our data support ATP-dependent activation of downstream defence activities, the precise molecular output of this synergy remains unresolved at the level of bulk DNA integrity measurements.

Genetic analysis further suggested that Gabija and Septu contribute to the synergy in different manners. Point mutations in the PtuB HNH endonuclease of Septu type I abolished synergy with Tmn, consistent with a model in which PtuA detects the low ATP state created by Tmn and PtuB executes the downstream Septu effector response (**Fig. S6c**). By contrast, point mutations in the GajA TOPRIM domain and in GajB catalytic residues did not abolish Gabija-Tmn synergy (**Fig. S6c**), even though these GajA substitutions disrupt Gabija defence in other contexts^21^. These findings suggest that the Gabija contribution to synergy under Tmn-activating conditions does not require the canonical TOPRIM-dependent DNA cleavage output of this system. In addition, GajB ATPase activity was not required although GajB itself was (**Fig. S6c**), consistent with a non-catalytic or complex-stabilising role for GajB in enabling Gabija function in synergy with Tmn^21,42,46^.

To elucidate how this layered defensive response reshapes infection outcomes at single-cell resolution, we used microfluidics. Co-expression of Tmn with either Gabija or Septu type I substantially increased the frequency of plasmolysis during SLUR435 infection but did not markedly shift the timing of the first lysis event (**Fig. 6c,d**, **Fig. S6d**). Because Tmn remains the principal driver of ATP depletion in these backgrounds (**Fig. 6b**), the increased plasmolysis frequency is unlikely to reflect stronger primary activation of Tmn. Instead, it is most consistent with downstream defence activity amplifying the physiological consequences of Tmn-induced ATP collapse, for example, by pushing more cells past the morphological threshold used to score plasmolysis and/or by stabilising or prolonging the plasmolysed state so it is more readily detected. In line with a shift towards poorly productive infection outcomes, lysis of Tmn+Gabija and Tmn+Septu strains was substantially less likely to seed secondary infections and rarely resulted in immediate trench clearance (**Fig. 6e,f**), consistent with strongly reduced release of infectious progeny. Thus, although the onset of lysis was largely unchanged, the combined systems shifted infection outcomes towards plasmolysis-associated, low-productivity events.

Together, these findings support a model in which Tmn occupies an upstream position in a layered antiviral cascade. Tmn translates phage infection into an early ATP-collapse signal that activates otherwise silent ATP-sensing defences, which then reshape the course of infection, suppressing phage maturation and secondary spread.

## Discussion

In this study, we characterised the Tmn defence system, a membrane integrated P-loop NTPase of the YobI family that protects bacteria against phage infection by inducing a distinct, in some cases, reversible, physiological state. Upon detecting infection via direct interaction with the replisome-associated phage protein RIIB, Tmn couples ATP hydrolysis to selective magnesium export, leading to plasmolysis and arrest of phage development (**Fig. 6g**). This response occurs without detectable membrane depolarisation or gross permeability increase, distinguishing Tmn from membrane-disruptive abortive infection mechanisms^17,47,48^. At the single-cell level, plasmolysis can be transient, and a subset of Tmn-expressing cells recover growth after infection, indicating that Tmn can impose a protective pause rather than deterministically committing cells to immediate death. Even when infected cells ultimately lyse, infection outcomes are shifted towards low phage production, limiting release of infectious progeny and thereby protecting neighbouring cells at the population level. Together, these findings suggest that Tmn-induced plasmolysis defines a window in which phage progression is curtailed sufficiently to permit occasional recovery and, more broadly, to suppress onward spread of the phage. Thus, Tmn serves both as an intracellular defence and a mechanism for population-level protection.

Cryo-EM analysis, supported by AlphaFold models, revealed that Tmn assembles into a membrane-anchored decamer with extended cytosolic arms. This decameric organisation is unusual among P-loop NTPases, which more commonly form hexamers^25^ or, in the case of Avs, tetramers, and provides a large, radially arranged scaffold that supports trigger engagement. The cytosolic domain of Tmn forms a solenoid-like array of α-helical repeats, with a distinct arrangement of HEAT- and ARM-like repeats^33^. This arm architecture appears to provide an interaction interface for phage-encoded triggers. Given the diversity of Tmn homologues across bacterial species, particularly in the cytosolic arms, some of these likely respond to alternative phage triggers or export different ions, suggestive of modular adaptations of the core Tmn mechanism.

Conceptually, Tmn expands the emerging set of membrane-integrated defences that combine sensing and effector functions within a single, large polypeptide. Whereas many prokaryotic STAND-based defences and abortive infection systems rely on rapid, often irreversible cellular damage^10,15,49^, Tmn enforces a strong but potentially reversible physiological constrain (plasmolysis) that arrests infection while preserving a route to recovery.

The timing and specificity of Tmn activation are central to its efficacy in phage defence. RIIB is expressed early in infection, and a single amino acid change (D219) abolishes Tmn activation, pointing to a fine-tuned interaction. Because RIIB is produced before late infection events that irreversibly damage the host, its detection provides Tmn with an early window to arrest phage progression before the takeover of the infected cell is complete. Notably, RIIB variants can remain detectably associated with Tmn yet fail to activate it, indicating that binding and activation can be uncoupled and supporting a productive engagement model in which specific contacts or geometries are required to activate the Tmn NTPase. This mechanism mirrors eukaryotic pattern-recognition pathways in which trigger molecule binding must be transduced into an activated conformational and/or oligomeric state to enable catalysis or signalling^50^. How Tmn defence is deactivated once phage development is abrogated, remains unclear. The ability of some cells to recover suggests that Tmn activation is reversible, potentially being regulated by turnover of RIIB, disassembly or conformational change of the Tmn complex, or changes in gene expression.

Tmn also serves as a metabolic trigger of cooperative defence involving multiple systems (**Fig. 6g**). By depleting intracellular ATP, Tmn can activate ATP-sensitive systems such as Gabija and Septu type I, which contain ATP-responsive components that are restrained under ATP-replete conditions but are activated as the ATP level drops^42,45,46^. At the single-cell level, the combinations of Tmn with Gabija and Septu increase the penetrance of plasmolysis without substantially shifting first-lysis timing, consistent with downstream defence activities enhancing and/or prolonging the plasmolysis-associated state after Tmn initiates ATP collapse. The combination strains also show reduced secondary infection and trench clearance, indicating that layered activation reshapes the course of infection towards low-progeny outcomes. Although the precise downstream effector outputs remain unresolved in this context, these findings support a view in which ATP depletion operates as a shared infection signal that coordinates distinct defence modules into a temporally ordered, synergistic antiviral cascade.

In summary, Tmn links detection of an early phage replisome factor (and possibly, other triggers) to ATP turnover and selective Mg^2+^ export, inducing a plasmolysis-associated antiviral state without detectable membrane depolarisation or gross leakage. The ability to separate RIIB association from activation indicates that trigger recognition is translated into defence through a discrete activation step, rather than through stable complex formation alone. By generating an early ATP-collapse signal that can engage otherwise silent ATP-sensing systems such as Gabija and Septu type I, Tmn provides a mechanistic basis for synergy in antiphage defence and illustrates how changes in cellular energy state can coordinate layered antiviral responses.

## Data availability

All data generated or analysed during this study are included in this published article and its supplementary information files. Raw sequencing reads have been deposited at Zenodo https://doi.org/10.5281/zenodo.18819000, and RNAseq data at Zenodo https://doi.org/10.5281/zenodo.18819242. The cryo-EM density map of the Tmn complex has been deposited in the Electron Microscopy Data Bank (EMDB) under accession code EMD-75267. No atomic coordinates were deposited because the map resolution was insufficient for atomic model building.

## Code availability

Custom code used for the phylogenetic and RNAseq analysis is available in the GitHub repository https://github.com/garushyants/tmn_phylo.

## Acknowledgements

We thank Alexander Harms (ETH Zurich) for kindly providing the BASEL collection of phages, and Andrew Millard (University of Leicester) for phage SLUR435. We also thank Victor Tobiasson, Harutyun Sahakyan, Pascal Mutz and Ryan Bell (National Institutes of Health, NIH) for discussions, and J. De La Cruz (MSK) for assistance with cryo-EM data collection. Y.U. was supported by an NIHR Southampton Biomedical Research Centre Postdoctoral Bridging Fellowship. F.N. was supported by a Wessex Health Partners and National Institute for Health and Care Research Wessex Experimental Medicine Network seed fund. S.K.G. and E.V.K. were supported by the Intramural Research Program of the National Library of Medicine, NIH. D.J.P. was supported by NIH grant GM145888, the Maloris Foundation, and the Memorial Sloan Kettering Cancer Center Core Grant (P30-CA008748). In addition to the MSKCC cryo-EM facilities, portions of this work were performed at the National Center for CryoEM Access and Training (NCCAT) and the Simons Electron Microscopy Center at the New York Structural Biology Center, supported by the NIH Common Fund Transformative High Resolution Cryo-Electron Microscopy Program (U24 GM129539), the Simons Foundation (SF349247), and the New York State Assembly. The contributions of the NIH authors are considered Works of the United States Government. The findings and conclusions presented in this paper are those of the authors and do not necessarily reflect the views of the NIH or the U.S. Department of Health and Human Services. The research in S.B. lab was supported by the Wellcome Trust Award (RG89305), a University Startup Award for Lectureship in Synthetic Biology (NKXY ISSF3/46), an EPSRC New Investigator Award (EP/W032813/1) and a seed fund from the School of Technology, University of Cambridge.

## Author contributions

FLN conceptualised the study. YW, ZZ, SKG, SB, RL, TA, DJP, EVK, and FLN designed the experiments. YW, ZZ, SKG, RD, JAM, MJC, TA, SB and RL performed the experiments. YW, ZZ, SKG, SB, RL, TA, and FLN analysed the data. SB, YEG, EVK and FLN funded the project. YW, ZZ, SKG, SB, DJP, RL, EVK and FLN wrote the paper, while all authors reviewed at least one draft.

## Competing interests

The authors declare no competing interests.

## Methods

### Bacteria and phages

*E. coli* strain Dh5α was used to clone plasmids pACYC-duet or pCOLA-duet with Tmn variants or phage proteins. *E. coli* BL21-AI cells containing plasmid(s) with Tmn, Tmn mutations, and phage proteins were used for phage assays. Bacterial strains were grown at 37 °C in Lysogeny Broth (LB) with 180 rpm shaking for liquid cultures, or in LB agar (LBA) plates for solid cultures. Strains containing plasmid pACYC-duet or pCOLA-duet were grown in media supplemented with 25 µg/ml of chloramphenicol or 100 µg/ml of kanamycin, respectively. Phages used in this study are described in **Supplementary Table 4**.

### Phylogenetic analysis of Tmn

Summary tables from the precomputed PADLOC database (DB) over RefSeq v209 were downloaded from the PADLOC DB webpage (https://padloc.otago.ac.nz/padloc/refseq/)^23,24^. All records containing Tmn were extracted, yielding 7,255 records from 6,844 genomes. To ensure high-quality data, records from genomes that had been removed from RefSeq as of January 28, 2024, were excluded. Additionally, we filtered out proteins covering less than 50% of the PADLOC Tmn HMM profile and with e-value ≥ 1e-10, as well as those marked as pseudogenes. This filtering resulted in a dataset of 6,059 proteins from 6,031 genomes. For further analysis, the dataset was dereplicated with MMseqs2 v17.b804f^51^ with the following parameters: -c 0.9 --min-seq-id 0.98 – yielding 859 unique proteins. These sequences were downloaded from NCBI Batch Entrez and aligned using MUSCLE5^52^ with the -align command. The resulting alignments were inspected manually and all incomplete sequences missing P-loop ATPase domain (60 cases) and those having inactivated Walker A (14 cases) were removed. The resulting final set of 785 proteins was realigned with MUSCLE5^52^ and trimmed with pytrimAl^53^ using the method="gappyout" parameter. The trimmed alignment was used as input to IQ-TREE v2.1.3^54,55^ for phylogenetic reconstruction with the parameters -B 1000 -mset WAG,LG. Clades within the phylogenetic tree were assigned using TreeCluster^56^ with the Med Clade method and a cutoff of 3.5. The final phylogenetic tree was visualized using the ggtree package in R.

### Identification of conserved residues

Residues essential for Tmn function were identified using the following metrics for each alignment column: Shannon entropy < 1 and conservation >90%, with no more than 20% gaps. Conservation scores for the alignment were calculated with PyCanal package (https://github.com/jafetgado/PyCanal). The Logos were visualized with Logomaker python package^57^.

### Characterisation of genomic contexts

For each protein in the dataset, one representative genome was selected, prioritising Complete and Chromosome level assemblies when available. Taxonomic assignments for selected genomes were obtained with NCBI Datasets command-line tools (v18.15.0). Chromosomal or plasmid localisation was determined using geNomad^58^ and Platon (v1.6)^59^ for plasmid prediction. Gene annotations were derived from PADLOC defence system annotations^23,24^, the PFAM-A database^60^, and the geNomad database (v1.9). Genomic contexts were visualized in R (v4.4.2) using ggplot2 (https://github.com/tidyverse/ggplot2) and gggenes (https://github.com/wilkox/gggenes).

### Cloning of Tmn variants, mutants, and phage proteins

Plasmids constructed and primers used in this work can be found in **Supplementary Tables 5** and **6**, respectively. Tmn from *E. coli* ECOR25, ECOR70, b1359, *P. aeruginosa* LN11 and *K. pneumonia* OXA-48 were cloned into pACYC-duet by Gibson assembly. Phage proteins were cloned into pCOLA-duet by Gibson assembly. The plasmids were recovered in Dh5α cells, extracted using NucleoSpin Plasmid QuickPure Kit (Thermo Fisher Scientific) and confirmed by sequencing at Eurofins Genomics or Plasmidsaurus (USA). Mutations of Tmn and phage proteins were engineered by around-the-horn PCR, and confirmed by Sanger sequencing at Eurofins Genomics. Plasmids were transformed individually or in combinations into competent BL21-AI cells prepared using the Mix&Go! *E. coli* Transformation Kit (Zymo).

### Efficiency of platting

Overnight cultures of the bacteria were diluted 1:50 in LB containing antibiotics, induced with 0.2% arabinose, and incubated for 4 hours before being used in double agar overlay assays. For this, bacterial cultures were mixed with 0.6% top agar and overlaid on LBA plates. Ten-fold serial dilutions of the phage stocks were spotted onto the bacterial lawn and the plates incubated overnight at 37 °C. The phage plaques were counted and used to calculate the EOP relative to the control. Statistical significance was determined using the multiple comparison function from Two-way ANOVA with a p-value of <0.01.

### Time post infection assay

Overnight bacterial cultures were diluted to an optical density at 600 nm of 0.1 in LB containing antibiotics and 0.2% arabinose. The cultures were infected with phage at an MOI of 10. At 0, 15, 30, 60 and 120-minutes post infection, a sample was taken and centrifuged at 12,000 × *g* for 2 minutes. The supernatant was serially diluted, and the phages were quantified by plaque assay on a bacterial lawn of cells with YFP. PFUs were counted after overnight incubation at 37 °C. To determine the CFUs, the pellets were washed twice with LB, the cells resuspended and serially diluted. The dilutions were spotted onto LBA plates supplemented with antibiotics, and the plates incubated overnight at 37 °C.

### Liquid assay

Overnight bacterial cultures were diluted to an optical density at 600 nm of 0.1 in LB containing antibiotics and 0.2% arabinose. The bacterial suspension was distributed into wells of a 96-well plate to which phage dilutions or LB were added. The plates were incubated in a Clariostar Plus plate reader at 37 °C with shaking at 200 rpm, with optical density at 600 nm measured every 10 min for 24 h.

### Generation of phage escape mutants

Phage plaques obtained in double layer agar plates with Tmn were picked and dissolved in 30 µl of LB. The recovered phages were produced in liquid culture using the Tmn-expressing strain as the host. The produced phages was serially diluted and spotted onto bacterial laws of Tmn-expressing and control cells, to identify those phages able to escape Tmn defence.

### Phage DNA extraction and sequencing

Phage DNA was extracted using the phenol-chloroform method. Briefly, DNase I and RNase were added to the phage stock at 1 µg/ml each, and the solution was incubated for 30 min. EDTA, proteinase K, and SDS were added to the solution at 20 mM, 50 µg/ml, and 0.5%, respectively, and incubated for 1 hour at 56 °C. An equal volume of phenol was added to the mixture, vortexed, and centrifuged at 3,000 × *g* for 10 min, at room temperature. The aqueous phase was recovered, mixed with an equal volume of 1:1 phenol:chloroform, and centrifuged. The procedure was repeated with chloroform. After centrifugation, the aqueous phase was recovered and mixed with 0.1 volumes of 3 M sodium acetate, pH 5.0 and 2.5 volumes of ice-cold absolute ethanol. The mixture was incubated at -20 °C for 1 hour and centrifuged at 14,000 × *g* for 15 min. The DNA pellet was resuspended in ice-cold 70% ethanol and centrifuged for 5 min. The pellet was let dry at room temperature and resuspended in nuclease-free water. The quality and quantity of extracted phage DNA were estimated using a Nanodrop and a Qubit fluorometer, respectively. Phage samples were sequenced at Novogene. Sample libraries were prepared using Novogene NGS DNA Library Prep Set (Cat. No. PT004), and sequenced on an Illumina platform, producing 1-2 Gb reads.

### Sequencing and analysis of phage escape mutants

For wild-type phage, seqtk version 1.3 (https://github.com/lh3/seqtk) was used to sample reads at 100x depth of the phage genomes. The subset reads were assembled using SPAdes version v3.15.3^61^ using default settings. For phage escape mutants, all reads were mapped to the assembled wild-type phage genome using bwa version 0.7.17^62^ and samtools version 1.11^63^. SNP calling was performed using gatk version 4.2.6.1^64^.

### RNAseq

Overnight cultures were diluted 1:100 in LB supplemented with antibiotics and incubated at 37 °C, 180 rpm for 2 hours. The cells were infected with phage at an MOI of 10. After 15 min of incubation, cells were centrifuged and the RNA of the cell pellet immediately extracted using the RNease Kit (Qiagen). RNA was sequenced at Novogene (Novaseq 6,000, 2 × 150 bp). Reads were trimmed with Fastp (v. 0.23.4)^65^, aligned with Rsubread^66^, and analysed with edgeR package v3 in R (https://bioconductor.org/packages/release/bioc/html/edgeR.html).

### Membrane permeability and depolarisation

Overnight cultures of control (YFP) and Tmn-expressing cells were diluted to an OD at 600 nm of 0.1 in LB supplemented with antibiotics and 0.2% arabinose. The bacterial cultures were incubated at 37 °C, 200 rpm, until and OD600 of 0.4. The OD of the cultures were adjusted to 0.05 in LB supplemented with antibiotics, 0.2% arabinose, and 1% DMSO. For membrane permeability assays, PI was added to a final concentration of 5 µg/ml, while for membrane depolarisation, DISC3(5) was added to a final concentration of 10 nM. The cultures were incubated at 37 °C, 200 rpm, for 1 hour, and phages were then added at an MOI of 10 or 0.01. OD600, DISC(3)5 (excitation: 600 nm; emission: 670 nm), and PI (excitation: 544 nm; emission: 612 nm) were measured every 10 min.

### Fluorescence microscopy

Overnight cultures of control (YFP) or Tmn-expressing cells were sub-cultured in LB with antibiotics and 0.2% arabinose for 2 hours. Agarose pads were prepared with 1% agarose in LB or LB supplemented with 1μg/ml of PI (for live/dead staining). For membrane staining, cells were washed with 1x PBS and resuspended with 0.1 mM FM4-64 (in PBS). Two μl of bacterial culture were seeded onto the agarose pads and covered with microscope cover slips prior to imaging. Confocal images and movies were acquired using a dual point-scanning Nikon A1R-si microscope equipped with a PInano Piezo stage (MCL), using a 60x PlanApo VC oil objective NA 1.40. Movies and images were acquired in galvanometer scanning mode using 488nm Laser (Coherent, 50mW), and a 561 nm Laser (Coherent, 50mW). Image processing and quantification was performed using FiJi software.

### Cross-sectional TEM

Overnight cultures of the bacteria were diluted 1:50 in LB containing antibiotics, induced with 0.2% arabinose, and incubated for 2 hours at 37 °C with shaking at 200 rpm. Phages were added at an MOI of 10 for the condition with phage infection. The cultures were incubated for 15 minutes at 37 °C with shaking at 200 rpm. Two mL of the cultures were centrifuged at 12,000 × *g* for 2 minutes and resuspended in 1 mL of 4% formalin, 3% glutaraldehyde and 0.1M PIPES. Bacterial cells were suspended in 5% sodium alginate and pelleted by centrifugation at 9000 rpm for 2 min. The resulting alginate–cell mixture was cross-linked by addition of 0.1 M calcium chloride and allowed to harden overnight to form calcium alginate pellets. Pellets were post fixed in 1% osmium tetroxide prepared in 0.1 M PIPES buffer, stained with 4% aqueous neodymium acetate, dehydrated through a graded ethanol series, infiltrated with the link reagent acetonitrile, and embedded in Spurr’s resin (Agar Scientific, Essex, UK). Ultrathin sections (70 nm) were cut using a Leica UC7 ultramicrotome, collected on 200 mesh Cu/Pd grids, stained with Reynolds’ lead citrate, and examined using an FEI Tecnai Spirit 12 transmission electron microscope equipped with a Morada G2 side mount camera. The total and cytoplasmic areas of at least 20 cells were calculated using Fiji^67^, and the relative cytoplasmic area was calculated as the ratio between cytoplasmic and total areas.

### Elemental analysis

Overnight bacterial cultures were diluted to an optical density at 600 nm of 0.1 in LB containing antibiotics and 0.2% arabinose. The cultures were infected with phage at an MOI of 10. At 0, 15, and 30-minutes post infection, a sample was taken and centrifuged at 12,000 × g for 2 minutes. Cell pellets were freeze-dried and weighed into Teflon vessels. Concentrated sub boiled HNO_3_ was added to the samples and then digested by microwave (CEM MARS6). The digested samples were evaporated to dryness and re-diluted in 3% HNO_3_ for analysis. Elemental analysis was carried out by ICP-OES (Thermo Scientific iCAP6500) using synthetic standards prepared from single element ICP-MS standards (Inorganic Ventures, Inc).

### Alkaline and native agarose gel electrophoresis

DNA integrity was assessed by native and alkaline agarose gel electrophoresis. For alkaline gels, agarose (0.8%, w/v) was prepared in 30 mM NaCl, 2 mM EDTA (pH 7.5), cast, and allowed to solidify. Before loading, gels were equilibrated for 1 h in alkaline electrophoresis buffer (30 mM NaOH, 2 mM EDTA). DNA samples were mixed with 6× alkaline loading buffer (180 mM NaOH, 6 mM EDTA, 18% glycerol, 0.05% bromophenol blue), heated at 70 °C for 5 min, and chilled on ice for 3 min before loading. Electrophoresis was performed in alkaline buffer at 3 V/cm until the dye front migrated approximately two-thirds of the gel length. Gels were then neutralised in 0.5 M Tris-HCl (pH 7.5) for 30 min and stained with SYBR Gold overnight prior to imaging. For comparison, the same samples were analysed by native agarose gel electrophoresis under non-denaturing conditions (1% agarose in 1× TAE), using standard loading dye and electrophoresis conditions, with SYBR Safe staining.

### Microfluidic device fabrication

Microfluidic devices were fabricated by standard PDMS soft lithography. Polydimethylsiloxane (PDMS, Dow Sylgard 184) base elastomer and curing agent were mixed at a ratio of 10:1 (w/w), degassed under vacuum for 30 min, poured over the SU-8 master, and degassed again to remove residual bubbles. PDMS was baked at 95 °C for 1 h, peeled off, and individual microfluidic devices were separated by cutting. Inlets and outlets were created in the individual devices using a 0.75 mm biopsy puncher. PDMS replicas were cleaned by sonication in isopropanol for 30 min, dried with compressed air, baked at 95 °C for 30 min, and sonicated again in deionised (DI) water for 30 min. Glass coverslips were cleaned by sonication in 1 M potassium hydroxide for 20 min, rinsed and sonicated in DI water for 20 min, dried with an air gun, and baked at 95 °C for 30 min. Devices were bonded (feature side to glass) using plasma treatment (2 min at 35 W and air pressure set to 0.2–0.3 mbar), followed by a post-bond bake at 95 °C for 30 min. Bonded devices were checked using a stereo microscope before cell loading for a timelapse imaging experiment.

### Single-cell phage infection assay and timelapse microscopy

Single-cell phage infection assays in microfluidic devices were performed as described previously^36^. *E. coli* cells carrying respective constructs were grown overnight from frozen stocks in LB Miller supplemented with relevant inducer, selection antibiotics, and surfactant Pluronic F-108 to minimise clumping and improve loading, then introduced into a microfluidic lane and allowed to diffuse into trenches before excess cells were removed by a short high-flowrate wash. To suppress inlet biofilm formation, device inlet was illuminated with 365-nm light for 10 min. The loaded cells were grown under constant supply of nutrients for several hours to reach steady-state exponential growth prior to phage infection. Infection was initiated by switching fresh growth medium to phage containing medium (prepared by diluting washed lysate to a target titre). Timelapse microscopy was performed on a Nikon Eclipse Ti2 inverted microscope equipped with a Hamamatsu C14440-20UP camera. Images were acquired using a 40×/0.95NA phase-contrast (PC) objective and a 1.5x post-objective magnifier. The microfluidic device was fixed on the motorised stage inside a temperature-controlled incubator maintained at 37 °C and continuously provided with media using a syringe pump connected via silicon tubing and 21-gauge needles. Multiple fields of view (FOVs) were imaged automatically using the automated stage, with focus stability maintained by Nikon’s Perfect Focus System to compensate for thermal drift. Images were collected every 2 min with 50 ms exposure time for PC images.

### Single-cell data analysis

Image data were recorded in ND2 format and converted into per-FOV image stacks (PNG snapshots) using custom Python scripts. Time series were registered to compensate for any XY stage drift, and each FOV was cropped into individual trench movies using a custom-designed image-processing pipeline as described previously^68^. Single cell segmentation was performed on the trench movies using Omnipose^69^, retrained on synthetic microfluidic trench datasets generated with virtual microscopy platform SyMBac^70^. A custom-designed lineage tracking program (https://github.com/erezli/MMLineageTracking) was used to extract features (e.g., cell size and fluorescence intensity) and track the cell lineages over time. Infection-related features were analysed using custom Python scripts.

### Protein expression and purification

The genes encoding Tmn-25 and T2 RIIB were cloned into the pET21a vector (Novagen) carrying ampicillin resistance and the pYB100 vector (NovoPro) carrying kanamycin resistance, respectively. Tmn was fused to a C-terminal Flag tag, and RIIB was fused to a C-terminal His6 tag. For recombinant expression in Expi293 cells, the genes encoding full-length Tmn or RIIB were codon-optimised and synthesised by Integrated DNA Technologies (IDT). The coding sequences were subsequently cloned into the pcDNA3.4 expression vector to generate constructs bearing either a C-terminal Flag tag or a C-terminal His₆ tag. All constructions were generated using Gibson assembly (NEB) and confirmed by sequencing.

The full-length Tmn membrane protein was overexpressed in C41 cells, a derivative of Escherichia coli BL21(DE3). Protein expression was induced with 0.5 mM IPTG (GoldBio) at 16 °C for 20 hours. Cells were harvested by centrifugation and resuspended in lysis buffer (25 mM Tris, pH 8.0, 500 mM NaCl, 2 mM β-mercaptoethanol, and cOmplete EDTA-free protease inhibitor). Cell lysis was performed by sonication, followed by the addition of 1% (w/v) Lauryl Maltose Neopentyl Glycol (LMNG, Anatrace) and 0.1% (w/v) cholesteryl hemisuccinate (CHS, Anatrace) to solubilize membrane proteins at 4 °C for 3 hours. Insoluble material was removed by centrifugation at 22,000 rpm for 1 hour using a JA-20 fixed-angle rotor (Avanti J-E series centrifuge, Beckman Coulter). The supernatant was incubated with anti-FLAG M2 affinity gel (Sigma, A2220) while rotating at 4 °C for 3 hours. The mixture was then loaded onto a gravity column and washed extensively with Wash Buffer I (25 mM Tris, pH 8.0, 500 mM NaCl, 2 mM β-mercaptoethanol, 0.1% (w/v) LMNG, and 0.01% (w/v) CHS), followed by Wash Buffer II (25 mM Tris, pH 8.0, 500 mM NaCl, 2 mM β-mercaptoethanol, 0.001% (w/v) LMNG, 0.0001% (w/v) CHS, and 0.00033% Glyco-diosgenin (GDN, Anatrace)). The protein was eluted with Wash Buffer II supplemented with 0.2 mg/ml 3× DYKDDDDK peptide (Pierce) while rotating at 4 °C for 30 minutes. The eluate was concentrated and further purified by size-exclusion chromatography (SEC) on a Superdex 6 Increase 10/300 GL column (GE Healthcare) equilibrated with SEC buffer (25 mM Tris, pH 8.0, 500 mM NaCl, 2 mM β-mercaptoethanol, 0.001% (w/v) LMNG, 0.0001% (w/v) CHS, and 0.00033% GDN). Fractions eluting at ∼11.6 ml, corresponding to high-order assemblies, were pooled, concentrated, and prepared for SDS-PAGE and cryo-EM analysis.

The Tmn-RIIB complex was co-expressed in C41 cells by co-transforming Tmn-Flag and RIIB-His constructs onto agar plates containing both ampicillin and kanamycin. A positive colony was picked and used for culture growth and protein expression. Protein expression was induced with 0.5 mM IPTG at 16 °C for 18 hours. Cell harvesting and complex purification were performed following the same procedure as described for Tmn alone.

For expression in Expi293 cells, Expi293F cells were cultured in Expi293 Expression Medium (Gibco) at 37 °C in a shaking incubator with 8% CO₂ and 70% relative humidity. When the cell density reached 3.0 × 10 cells/mL, plasmid DNA encoding full-length Tmn was mixed with 40-kDa linear polyethylenimine (PEI; Polysciences) at a DNA: PEI mass ratio of 1:2 and incubated for 30 min prior to transfection. Sodium butyrate was added to a final concentration of 10 mM, 16 h after transfection, to enhance protein expression. Cells were then cultured for an additional 3 days before harvest. To produce the Tmn-RIIB complex, plasmids encoding Tmn and RIIB were co-transfected at a 1:2 ratio under identical conditions. Cells were harvested 3 days after sodium butyrate addition by centrifugation at 2,500 × *g* for 20 min at 4 °C. Cell lysis and subsequent purification procedures followed the same protocols used for Tmn or the Tmn-RIIB complex expressed in *E. coli* C41 cells.

### Cryo-EM sample preparation and data collection

Purified full-length Tmn was concentrated to 6 mg/ml, and 0.5 mM FOM (fluorinated octyl maltoside) was added to reduce preferred orientation bias. A 4 µl aliquot of the sample was applied to glow-discharged holey gold grids (UltrAuFoil 300 mesh R1.2/1.3). The grids were blotted for 3.5 s with a blot force of 0 at 6 °C and 100% humidity and then plunge-frozen in liquid ethane using a Vitrobot Mark IV (FEI). Cryo-EM data were collected at the Memorial Sloan Kettering Cancer Center (MSKCC) on a Titan Krios G4 transmission electron microscope (FEI) operated at 300 kV, equipped with a Falcon 4i direct electron detector and controlled by EPU software. Movies were recorded in counting mode with a total electron dose of 60.3 e⁻/Å², a defocus range of -0.8 to -2.2 µm, and a physical pixel size of 0.725 Å.

### Cryo-EM data processing

A total of 7,157 movies were collected and processed using cryoSPARC^71^. Patch motion correction and patch contrast transfer function (CTF) estimation were applied to correct for beam-induced motion and to estimate CTF parameters, respectively. Micrographs with ice contamination, high astigmatism, or poor CTF fit resolution were excluded by applying appropriate threshold ranges in the *Manually Curate Exposures* job. High-quality micrographs were processed using the *Blob Picker* and *Extract* jobs, followed by 2D classification to generate templates for the *Template Picker*. Prior to 2D classification, the *Remove Duplicate Particles* job was used to eliminate overlapping or duplicate particles. Multiple rounds of 2D classification were then performed to discard junk particles, resulting in 10,427 high-quality particles for subsequent 3D reconstruction. *Ab initio* model reconstruction and heterogeneous refinement identified one volume with C1 symmetry corresponding to the Tmn decamer, using 4,025 particles. This volume was further refined through homogeneous refinement with C1 symmetry and non-uniform refinement with C5 symmetry, yielding a final resolution of 11.6 Å based on the Fourier shell correlation (FSC) 0.143 cutoff criterion.

### Model building, structure refinement, and visualisation

The Tmn decamer models were predicted using AlphaFold3^40^. Specifically, the Tmn N-terminal domain (residues 1-440) was predicted as a decamer (ipTM: 0.58; pTM: 0.60), whereas the full-length Tmn was predicted as a dimer (ipTM: 0.48; pTM: 0.50) due to size limitations in AF3 predictions. The full-length decamer structure was subsequently generated by superimposing the transmembrane domains, adjusting the short-leg regions, and assembling the complex in Chimera^72^. The model was further refined in COOT^73^ and PHENIX^74^, and all figures were visualised using ChimeraX^75^ and finalized in Adobe Photoshop. Structural alignment of cytoplasmic arms was performed with Icarus^31^ with –iterative parameter.

### Co-immunoprecipitation assay

To assess interactions between Tmn and RIIB, co-immunoprecipitation (Co-IP) assays were performed using Flag-tagged full-length Tmn (Tmn-FL) or Tmn truncation constructs in combination with His-tagged RIIB. Tmn truncations were generated using Gibson assembly and cloned into the same expression vector as Tmn-FL. All constructs were verified by Sanger sequencing prior to use. Co-transformation, protein expression, and cell culture conditions for Tmn-RIIB complexes were identical to those described above for Tmn-FL-RIIB expression. Protein purification was carried out as described above using Flag affinity resin in the presence of detergents, Mg²⁺ and ATPγS. Briefly, clarified cell lysates containing 0.01% (w/v) lauryl maltose neopentyl glycol (LMNG) and 0.001% (w/v) cholesteryl hemisuccinate (CHS) were incubated with anti-Flag resin for 2 h at 4 °C. The resin was washed extensively with 60 column volumes of wash buffer containing 25 mM HEPES (pH 7.5), 150 mM NaCl, 0.01% (w/v) LMNG, 0.001% (w/v) CHS, 5mM Mg^2+^, and 1 mM ATPγS.

Bound proteins were eluted using wash buffer supplemented with 0.2 mg/mL 3×FLAG peptide. Eluted samples were resolved by SDS-PAGE and transferred onto polyvinylidene fluoride (PVDF) membranes using a semi-dry transfer system (Bio-Rad). Membranes were blocked in 5% (w/v) non-fat dry milk (Bio-Rad) for 1 h at room temperature and incubated overnight at 4 °C with horseradish peroxidase (HRP)-conjugated anti-Flag (86861S, Cell Signaling Technology) or anti-His (MA1-21315-HRP, Thermo Fisher Scientific) antibodies at a 1:2,000 dilution (v/v). Following washing with 0.05% (v/v) Tween-20 in Tris-buffered saline (TBS), signals were visualised using enhanced chemiluminescence (Thermo Scientific) and quantified by densitometry using ImageJ 1.54g^76^.

### Blue-native PAGE

Purified Tmn-FL-RIIB complexes expressed in *E. coli* C41 cells (3 µg total protein) were analysed by native PAGE using 4-16% gradient gels (Thermo Scientific). Electrophoresis was performed at 4 °C at a constant voltage of 150 V for 60 min, followed by 250 V for an additional 60 min. Gels were fixed in 40% (v/v) methanol and 10% (v/v) acetic acid and briefly heated for 45 s, followed by shaking at room temperature for 15 min. Destaining was performed using 8% (v/v) acetic acid with a brief heating step. Bands were visualised using a Gel Doc XR+ imaging system (Bio-Rad) once an appropriate background level was achieved.

### Mass photometry

Mass photometry measurements were carried out using a Refeyn TwoMP instrument. A pre- assembled six-well sample cassette (Refeyn) was mounted onto a clean glass coverslip. For each measurement, 15 µL of freshly prepared SEC buffer (25 mM HEPES pH 7.5, 150 mM NaCl, 0.001% LMNG, 0.0001% CHS, and 0.00033% glyco-diosgenin (GDN)) was added to the well to establish the focal plane using the total internal reflection-based autofocus system. Purified Tmn-FL-RIIB complexes were diluted to 200 nM in SEC buffer, and 3 µL of the diluted sample was added to the buffer drop to achieve a final concentration of 33.3 nM. After stabilisation of the autofocus, movies were recorded for 60 s. Data were acquired using Refeyn AcquireMP (v2024.1.1.0) and analysed using Refeyn DiscoverMP (v2024.1.0.0). Contrast-to-mass calibration was performed using bovine serum albumin (BSA; Sigma), containing monomeric (66.5 kDa) and dimeric (132 kDa) species. Molecular mass distributions were analysed by Gaussian fitting within DiscoverMP to determine average molecular masses. Graphs were generated using GraphPad Prism 10.

### Statistical analysis

Unless stated otherwise, experimental data are presented as the mean of biological triplicates ± standard deviation. Statistical tests were performed using GraphPad Prism 10 and one sample t test or one-way ANOVA test.

---

**Figure S1.**
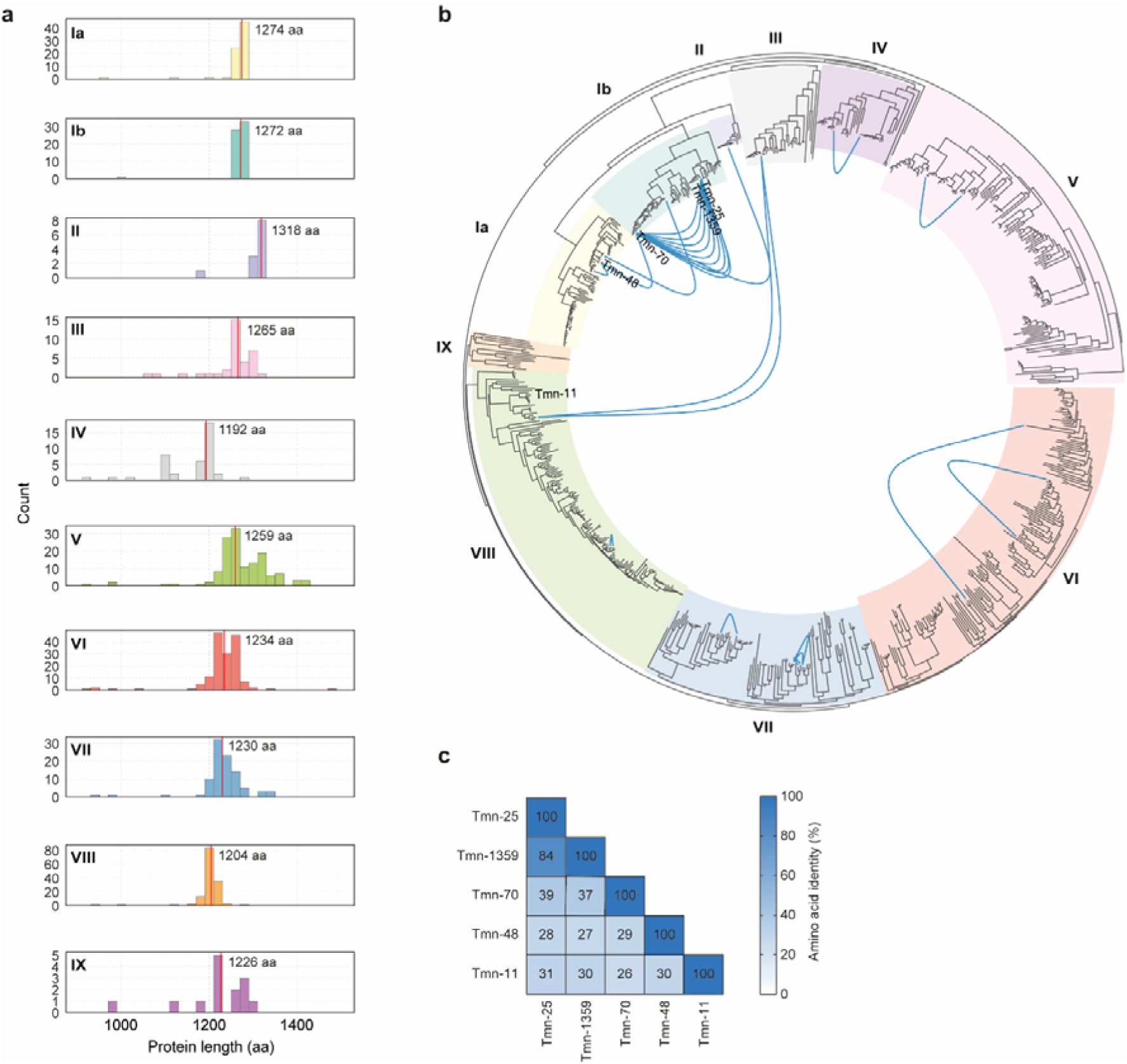
Length variation, co-occurrence, and sequence divergence across Tmn clades. **a**, Protein length distribution within Tmn clades. Histograms display the distribution of protein lengths for each clade, with the median lengths marked with a red line. **b,** Co-occurrence of two Tmn homologues within single genomes. Co-occurring proteins are indicated on the phylogeny and connected by blue lines. **c,** Pairwise amino acid identity (%) between the tested Tmn proteins.

**Figure S2.**
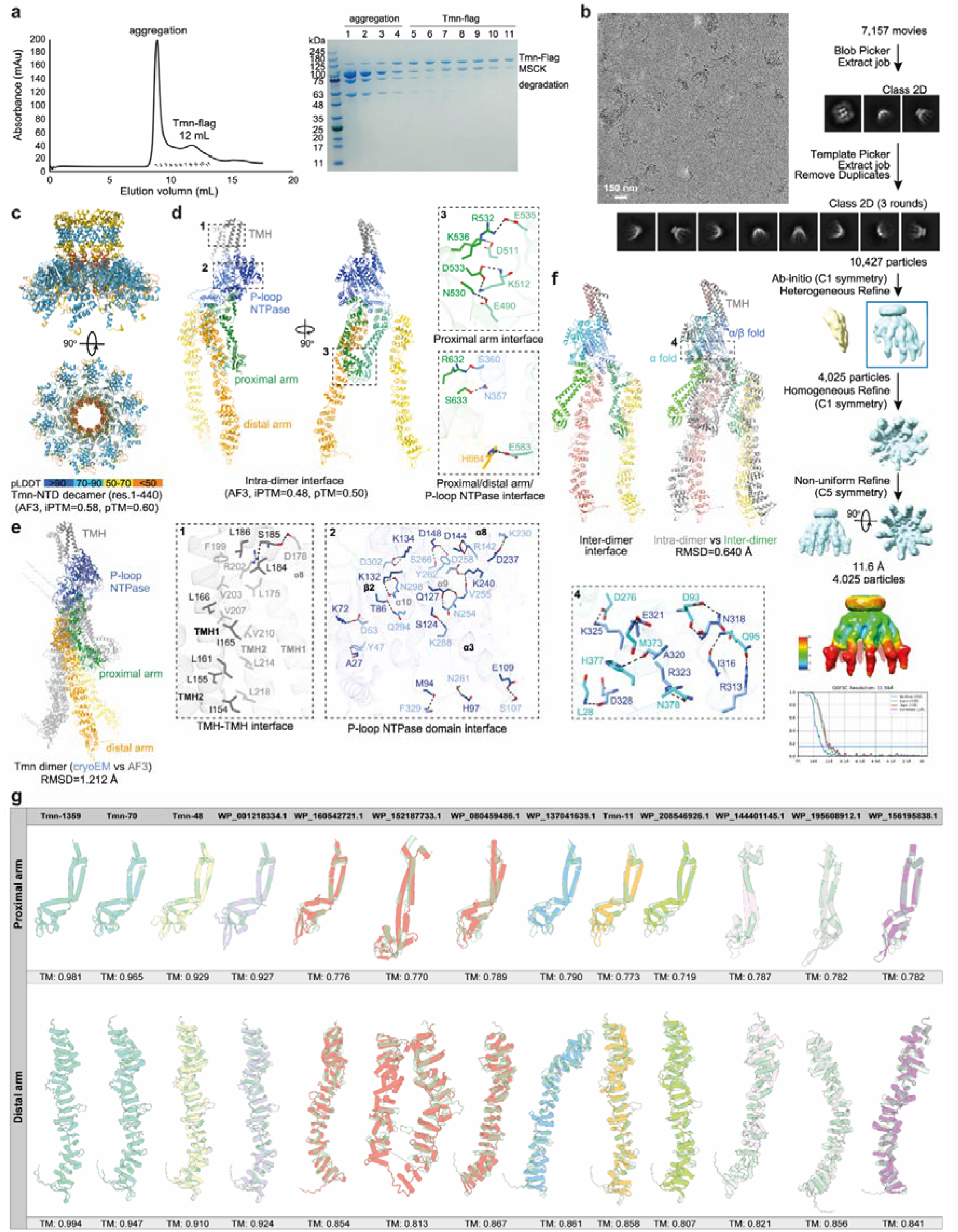
Structural analysis of Tmn. **a**, Size-exclusion chromatography (SEC) of purified Flag-tagged Tmn-25 using a Superose Increase 6 column. The major peak at 12 ml corresponds to decameric Tmn visualised by cryo-EM in panel (b), whereas earlier-eluting fractions indicate aggregate species. Corresponding SDS-PAGE gel analysis of selected fractions shows the presence of Tmn-Flag, minor contamination by mechanosensitive channel protein MscK, and partial degradation of Tmn-Flag, which is predominantly observed in the aggregate fractions. **b**, Representative cryo-EM raw micrograph of purified Tmn-25 particles, illustrating the heterogeneity and variation in particle orientation that limited high-resolution reconstruction. Overview of the cryo-EM image processing workflow for Tmn-25. 2D class averages and 3D reconstructions confirm a decameric structure. Bottom, local resolution map of the final Tmn reconstruction and gold-standard Fourier shell correlation (FSC) curve, indicating an overall resolution of ∼11.6 Å. **c**, AF3-predicted decameric assembly of the Tmn N-terminal region (residues 1-440) shown in side and top views. The model recapitulates the fivefold symmetric organisation and overall dimensions of the experimentally observed decamer, indicating the placement of the N-terminal region as a major contributor to the decameric core. **d**, Intra-dimer interface within the Tmn decamer. Two protomers forming a dimer are shown in distinct colours, highlighting contacts between (1) the TMH, (2) the P-loop NTPase domain, and (3) the proximal arm. Insets highlight representative contacts of boxed regions, with interface residues shown as sticks. **e**, Comparison of cryo-EM and AF3-predicted (ipTM=0.48, pTM=0.5) Tmn dimers, revealing similar overall conformations (RMSD = 1.212), except for the relative positions of the proximal and distal arm regions. **f**, Inter-dimer arrangements within the Tmn decamer (left) and its comparison with the intra-dimer arrangement (right). Superposition reveals similar overall conformations (RMSD = 0.640 Å), particularly within the Tmn core and the α/β fold of the P-loop NTPase domain. Some movements are observed in the α-helical regions of the P-loop NTPase domain, as well as in the proximal and distal arm regions. Inset shows modelled inter-dimer interfaces between neighbouring dimers within the decamer, highlighting contact surfaces involving the P-loop NTPase domains and adjacent arm segments that contribute to higher-order assembly. **g**, Structural comparison of proximal and distal arm regions across representative Tmn homologues from ten clades, superimposed against Tmn-25, illustrating conserved overall architecture with clade-dependent variation in arm length and curvature (clade colours are the same as in Figure 1c). TM values are provided in comparison with Tmn-25.

**Figure S3.**
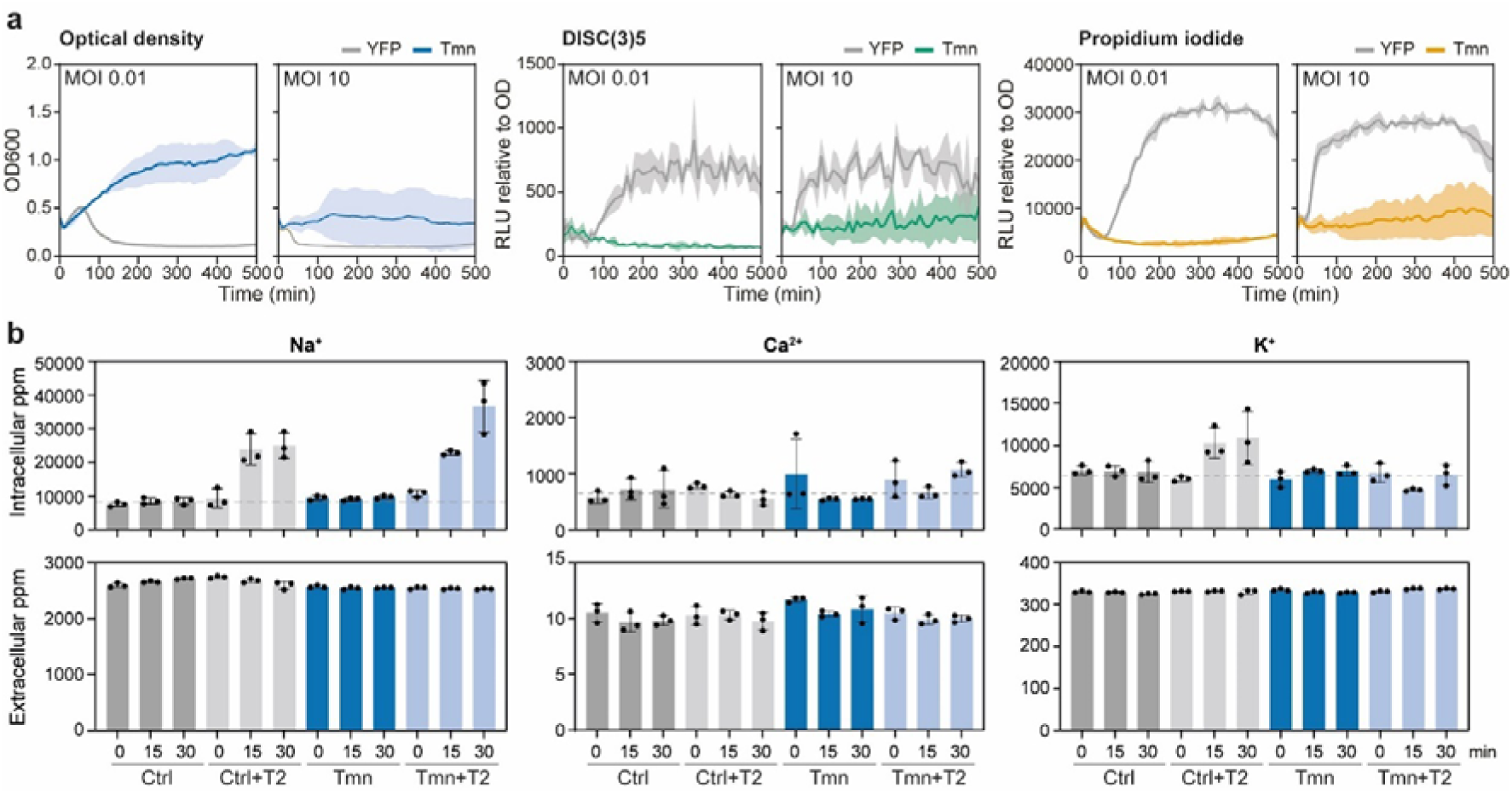
Tmn does not compromise membrane integrity or cause major ion leakage. **a**, Effect of phage infection at low (0.01) and high (10) multiplicity of infection (MOI) on control and Tmn cell growth (OD600), membrane depolarisation (DISC(3)5), and membrane permeability (propidium iodide, PI), measured over the course of 500 minutes. **b**, Quantification of intracellular (top) and extracellular (bottom) sodium (Na^+^), calcium (Ca^2+^), and potassium (K^+^) levels measured over time by electrospray ionisation mass spectrometry in uninfected and infected control and Tmn cells.

**Figure S4.**
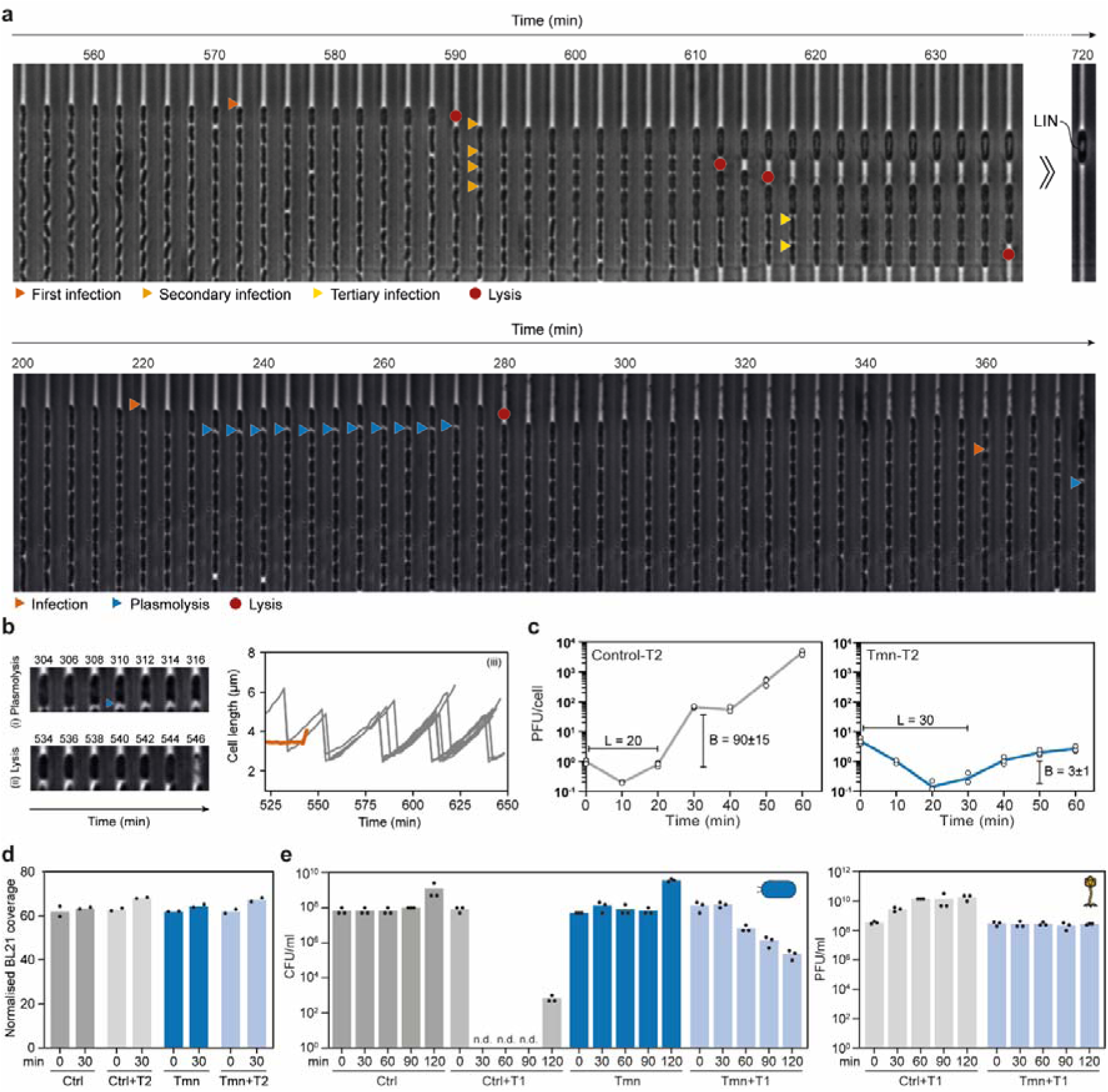
Tmn-induced plasmolysis delays lysis and prevents phage transmission. **a**, Time-lapse microscopy showing effective T2 infection and transmission in control cells (top) versus blocked transmission in Tmn-expressing cells (bottom). Initial infections (red) lead to secondary and tertiary infections in control cells but are halted in Tmn cells, which undergo plasmolysis (blue arrows). LIN, lysis inhibition. **b**, Zoomed examples of plasmolysis (i), lysis (ii), and successful sibling growth (iii) from Fig. 4b. **c**, One step growth curve of T2 in control (left) and Tmn-expressing (right) cells. B, burst size; L, latency period. **d**, Host genome integrity during infection measured by sequencing coverage. Bar plot shows the normalised breadth of coverage of the *E. coli* BL21 genome (fraction of genomic positions covered by mapped reads) at 0 and 30 min for T2 infected and uninfected control (Ctrl) and Tmn-expressing cells. Dots indicate independent biological replicates, and bars show the mean. **e**, Time post infection assays measuring bacterial (CFU/ml) and phage (PFU/ml) concentrations for T1 infected and uninfected control and Tmn cells. n.d., not detectable.

**Figure S5.**
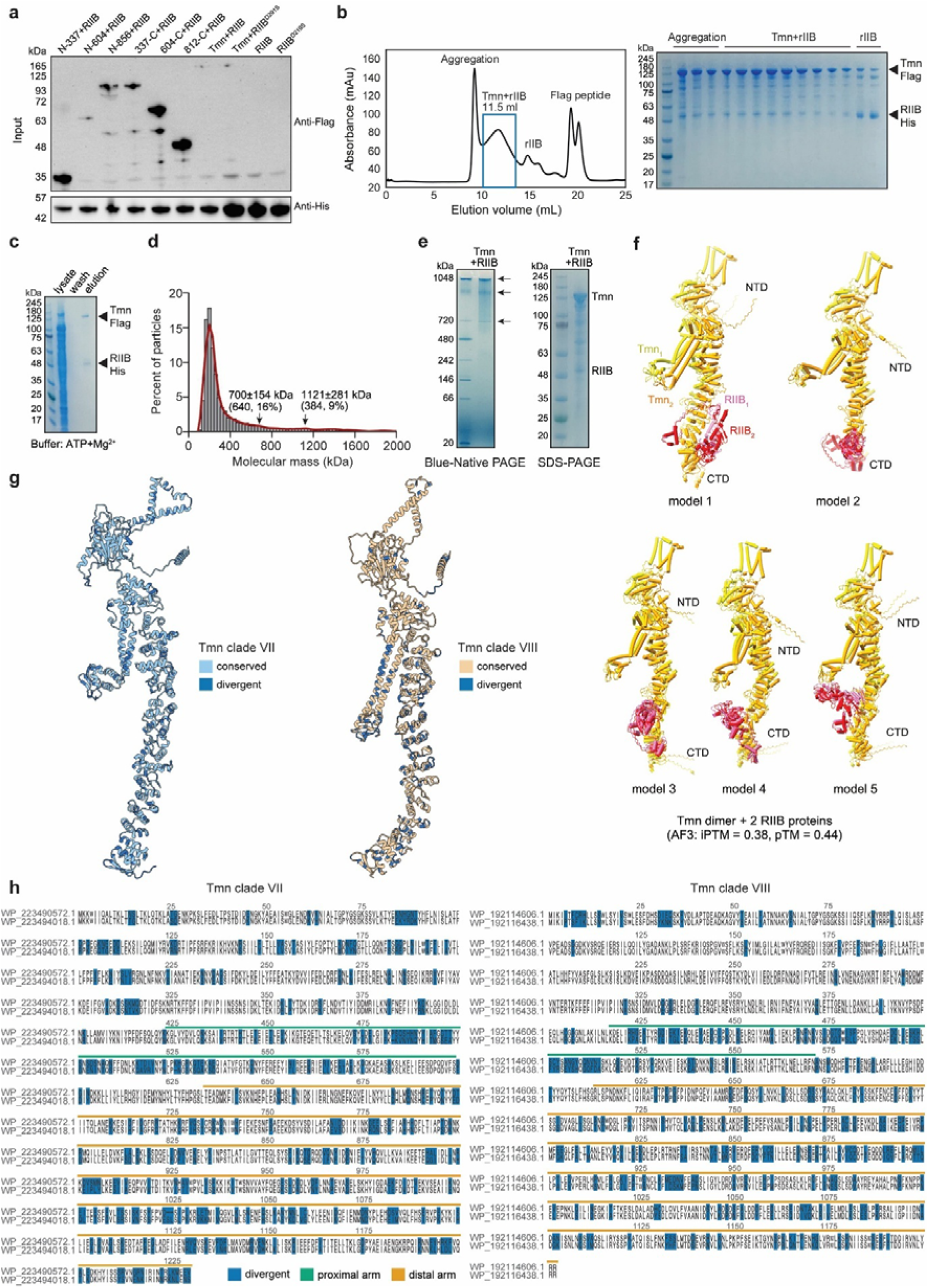
Biochemical characterisation of the Tmn-RIIB interaction and RIIB variants. **a**, Input immunoblots corresponding to the co-IP experiments in Fig. 5d, probed with anti-Flag (Tmn constructs) and anti-His (RIIB), indicating relative expression levels across constructs and conditions. **b,** Size exclusion chromatography (SEC) of co-expressed Tmn-25 and RIIB in *E*. *coli* C41 cells. Right: SDS-PAGE of SEC fractions. **c**, SDS-PAGE analysis of Tmn-Flag and RIIB-His co-expressed in *E*. *coli* C41 cells in the presence of 5 mM ATP and 10 mM Mg^2+^. **d**, Mass photometry of purified Tmn-RIIB samples showing heterogeneous particle populations, with major distribution peaks annotated on the plot. The initial large peak is buffer background. **e**, Blue native PAGE and SDS-PAGE analysis illustrating heterogeneous assemblies and co-recovery of both proteins under these conditions. **f**, AlphaFold 3 (AF3) co-folding predictions for a Tmn-25 dimer with two RIIB monomers (five independent models shown). Confidence metrics (ipTM and pTM) are indicated. **g**, Structural comparison of two pairs of closely related Tmn homologues encoded in the same genome, highlighting clusters of substitutions mapped onto AF3-predicted structures. Left, comparison of two clade VII Tmn variants from *Sutcliffiella horikoshii* (GCF_019931755.1), with residues that differ between the proteins highlighted in dark blue on the AF3 predicted structure of WP_223490572.1. Right, comparison of two clade VIII Tmn variants from *Shewanella sp.* BF02_Schw (GCF_014596695.2), with residues that differ between the proteins highlighted in dark blue on the AF3 predicted structure of WP_192114606.1. **h**, Pairwise sequence alignment of the Tmn homologue pairs shown in (g), with residues that differ between the two variants highlighted in blue.

**Figure S6.**
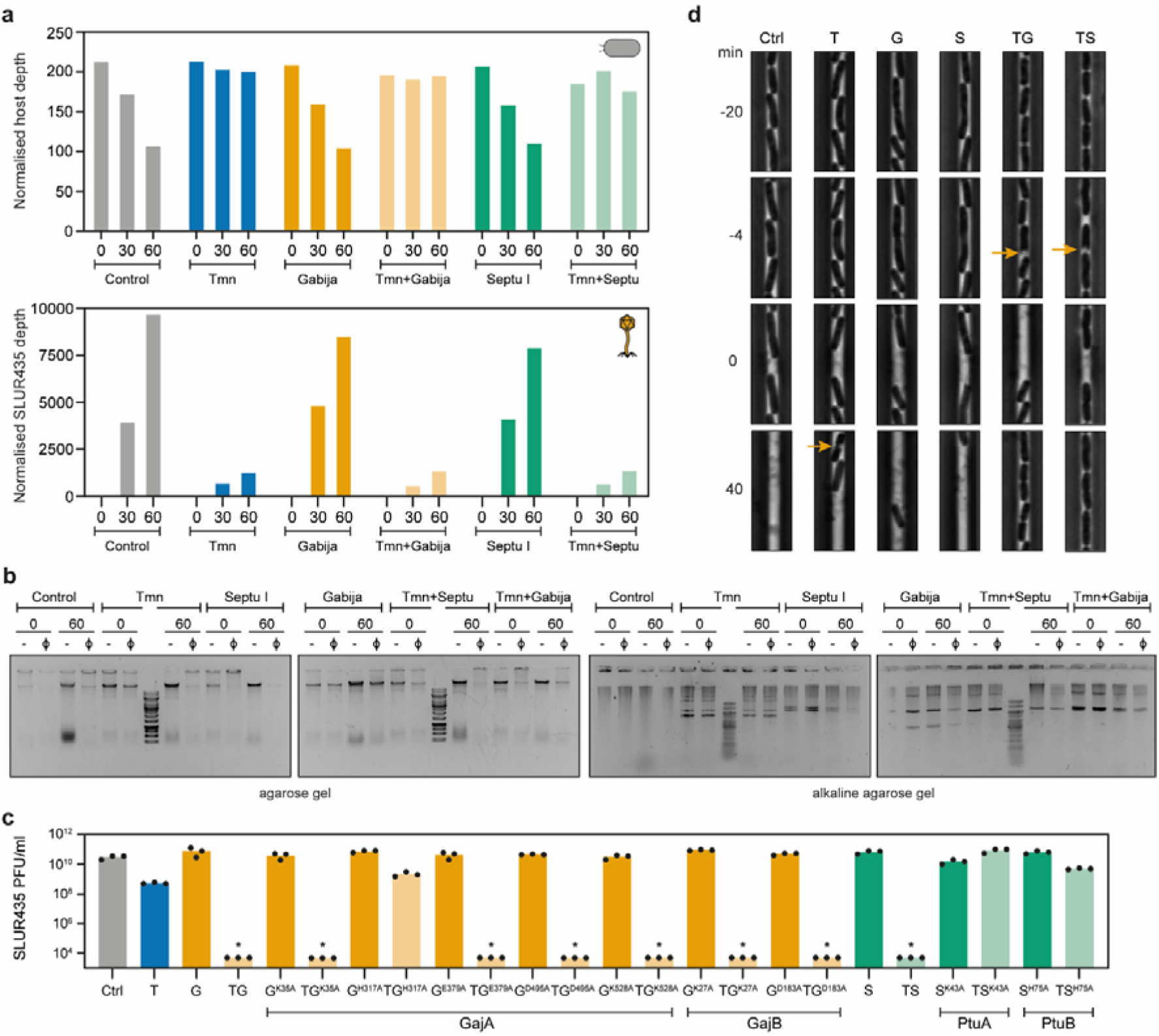
Genetic and single-cell dissection of Tmn synergy with Gabija and Septu type I. **a**, Quantification of bacterial host (top) and phage SLUR435 (bottom) DNA abundance during infection by Illumina sequencing. Coverage (depth) values are shown. **b**, Native (left) and alkaline (right) agarose analysis of DNA integrity during SLUR435 infection in strains expressing Tmn alone or in combination with Gabija/Septu. **c**, Mutational analysis of synergy between Tmn (T) and Gabija (G) or Septu (S) against phage SLUR435, measured as endpoint phage titre (PFU/ml). Dots indicate biological replicates; bars indicate mean. Asterisks denote significant differences (p < 0.05) relative to Tmn. **d,** Representative microfluidic trench trajectories illustrating infection-associated plasmolysis across strains. Example trenches are shown for control (Ctrl), Tmn (T), Gabija (G), Septu type I (S), Tmn+Gabija (TG), and Tmn+Septu (TS). Time is indicated relative to the first detectable lysis event (0 min). Arrows indicate plasmolysis events.

## Notes

### Competing Interest Statement

The authors have declared no competing interest.

