## Supplementary material for "Tmn blocks phage spread via plasmolysis and triggers synergistic defence responses": Fig. S

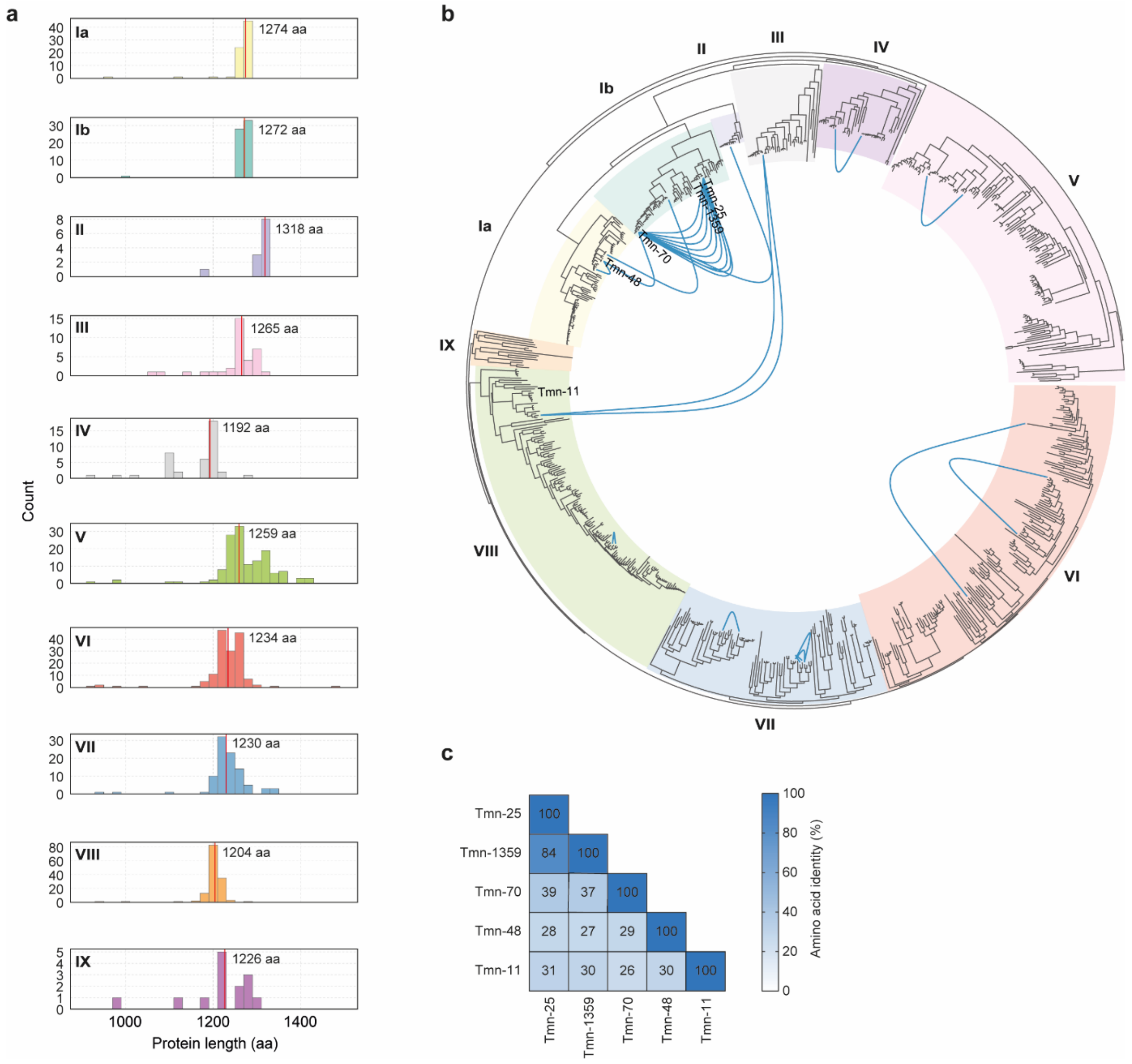


**Figure S1. Length variation, co-occurrence, and sequence divergence across Tmn clades. a,** Protein length distribution within Tmn clades. Histograms display the distribution of protein lengths for each clade, with the median lengths marked with a red line. **b,** Co-occurrence of two Tmn homologues within single genomes. Co-occurring proteins are indicated on the phylogeny and connected by blue lines. **c,** Pairwise amino acid identity (%) between the tested Tmn proteins.

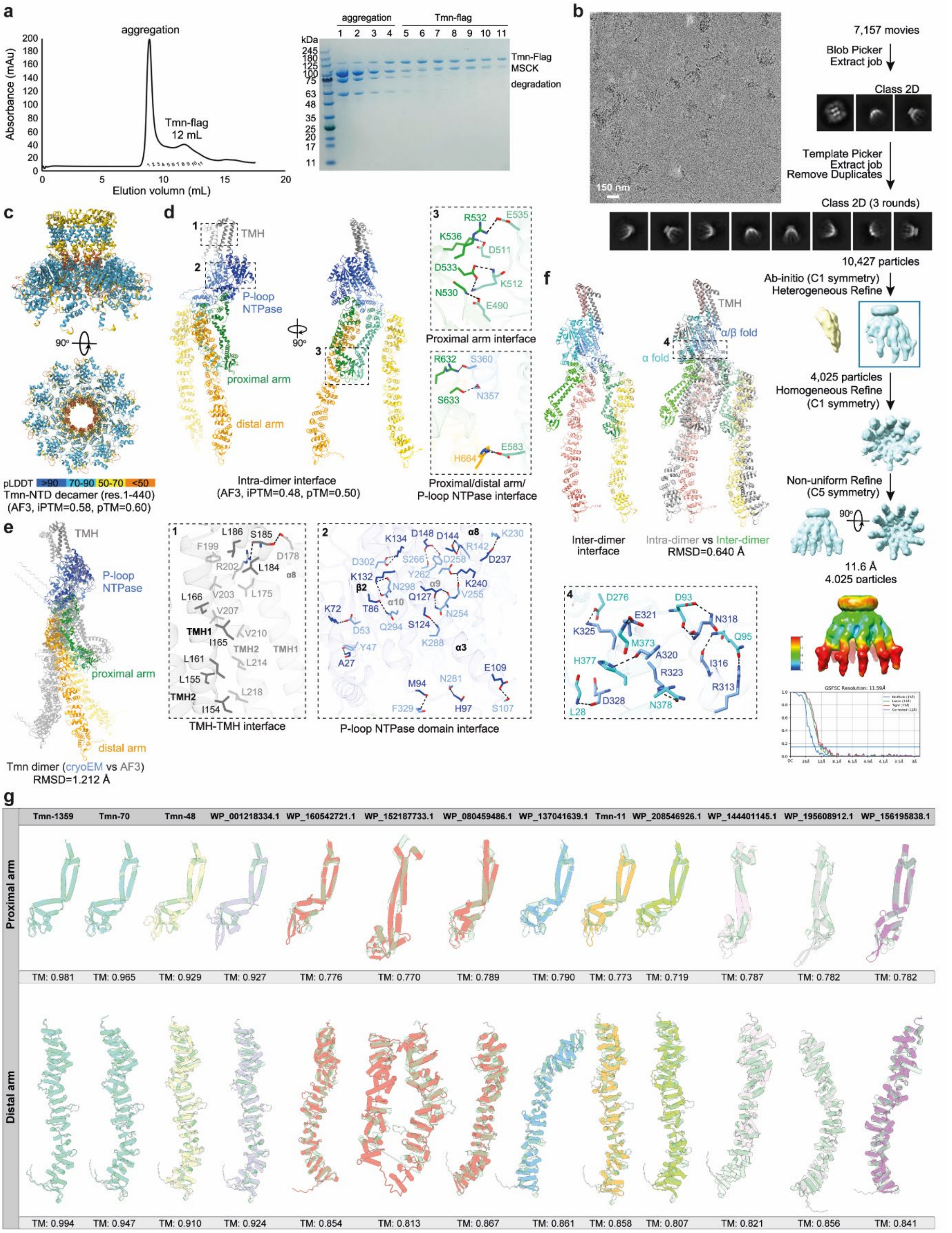


**Figure S2. Structural analysis of Tmn. a**, Size-exclusion chromatography (SEC) of purified Flag-tagged Tmn-25 using a Superose Increase 6 column. The major peak at 12 ml corresponds to decameric Tmn visualised by cryo-EM in panel (b), whereas earlier-eluting fractions indicate aggregate species. Corresponding SDS-PAGE gel analysis of selected fractions shows the presence of Tmn-Flag, minor contamination by mechanosensitive channel protein MscK, and partial degradation of Tmn-Flag, which is predominantly observed in the aggregate fractions. **b**, Representative cryo-EM raw micrograph of purified Tmn-25 particles, illustrating the heterogeneity and variation in particle orientation that limited high-resolution reconstruction. Overview of the cryo-EM image processing workflow for Tmn-25. 2D class averages and 3D reconstructions confirm a decameric structure. Bottom, local resolution map of the final Tmn reconstruction and gold-standard Fourier shell correlation (FSC) curve, indicating an overall resolution of ~11.6 Å. **c**, AF3-predicted decameric assembly of the Tmn N-terminal region (residues 1-440) shown in side and top views. The model recapitulates the fivefold symmetric organisation and overall dimensions of the experimentally observed decamer, indicating the placement of the N-terminal region as a major contributor to the decameric core. **d**, Intra-dimer interface within the Tmn decamer. Two protomers forming a dimer are shown in distinct colours, highlighting contacts between (1) the TMH, (2) the P-loop NTPase domain, and (3) the proximal arm. Insets highlight representative contacts of boxed regions, with interface residues shown as sticks. **e**, Comparison of cryo-EM and AF3-predicted (ipTM=0.48, pTM=0.5) Tmn dimers, revealing similar overall conformations (RMSD = 1.212), except for the relative positions of the proximal and distal arm regions. **f**, Inter-dimer arrangements within the Tmn decamer (left) and its comparison with the intra-dimer arrangement (right). Superposition reveals similar overall conformations (RMSD = 0.640 Å), particularly within the Tmn core and the α/β fold of the P-loop NTPase domain. Some movements are observed in the α-helical regions of the P-loop NTPase domain, as well as in the proximal and distal arm regions. Inset shows modelled inter-dimer interfaces between neighbouring dimers within the decamer, highlighting contact surfaces involving the P-loop NTPase domains and adjacent arm segments that contribute to higher-order assembly. **g**, Structural comparison of proximal and distal arm regions across representative Tmn homologues from ten clades, superimposed against Tmn-25, illustrating conserved overall architecture with clade-dependent variation in arm length and curvature (clade colours are the same as in Figure 1c). TM values are provided in comparison with Tmn-25.

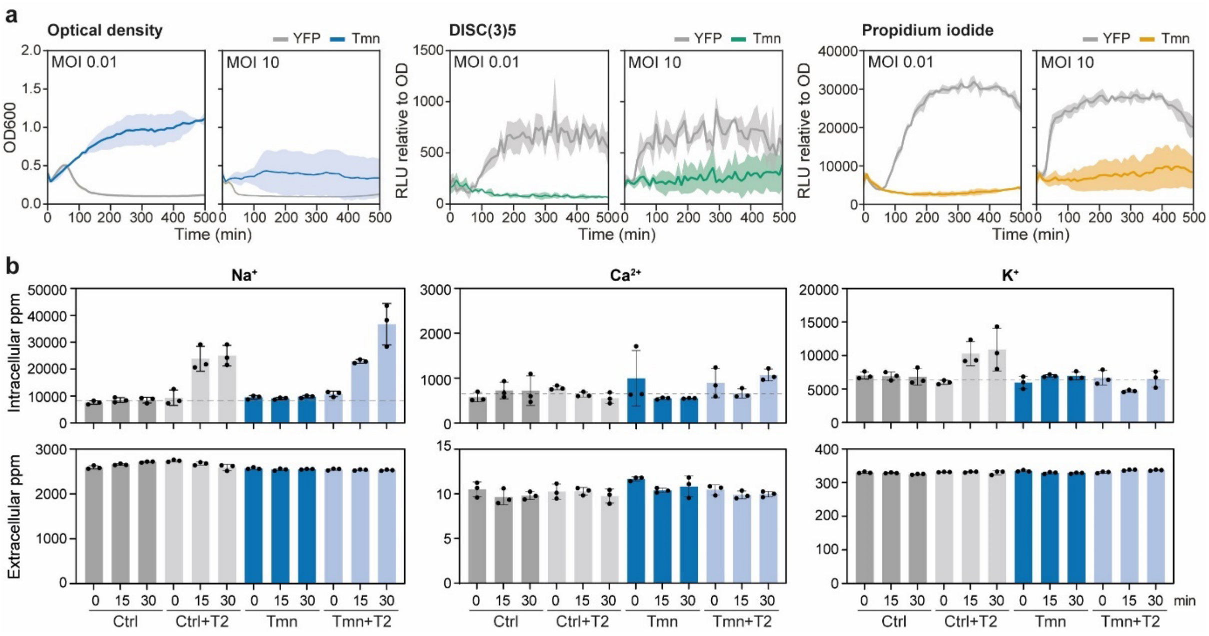


**Figure S3. Tmn does not compromise membrane integrity or cause major ion leakage. a,** Effect of phage infection at low (0.01) and high (10) multiplicity of infection (MOI) on control and Tmn cell growth (OD600), membrane depolarisation (DISC(3)5), and membrane permeability (propidium iodide, PI), measured over the course of 500 minutes. **b**, Quantification of intracellular (top) and extracellular (bottom) sodium ($Na^+$), calcium ($Ca^{2+}$), and potassium ($K^+$) levels measured over time by electrospray ionisation mass spectrometry in uninfected and infected control and Tmn cells.

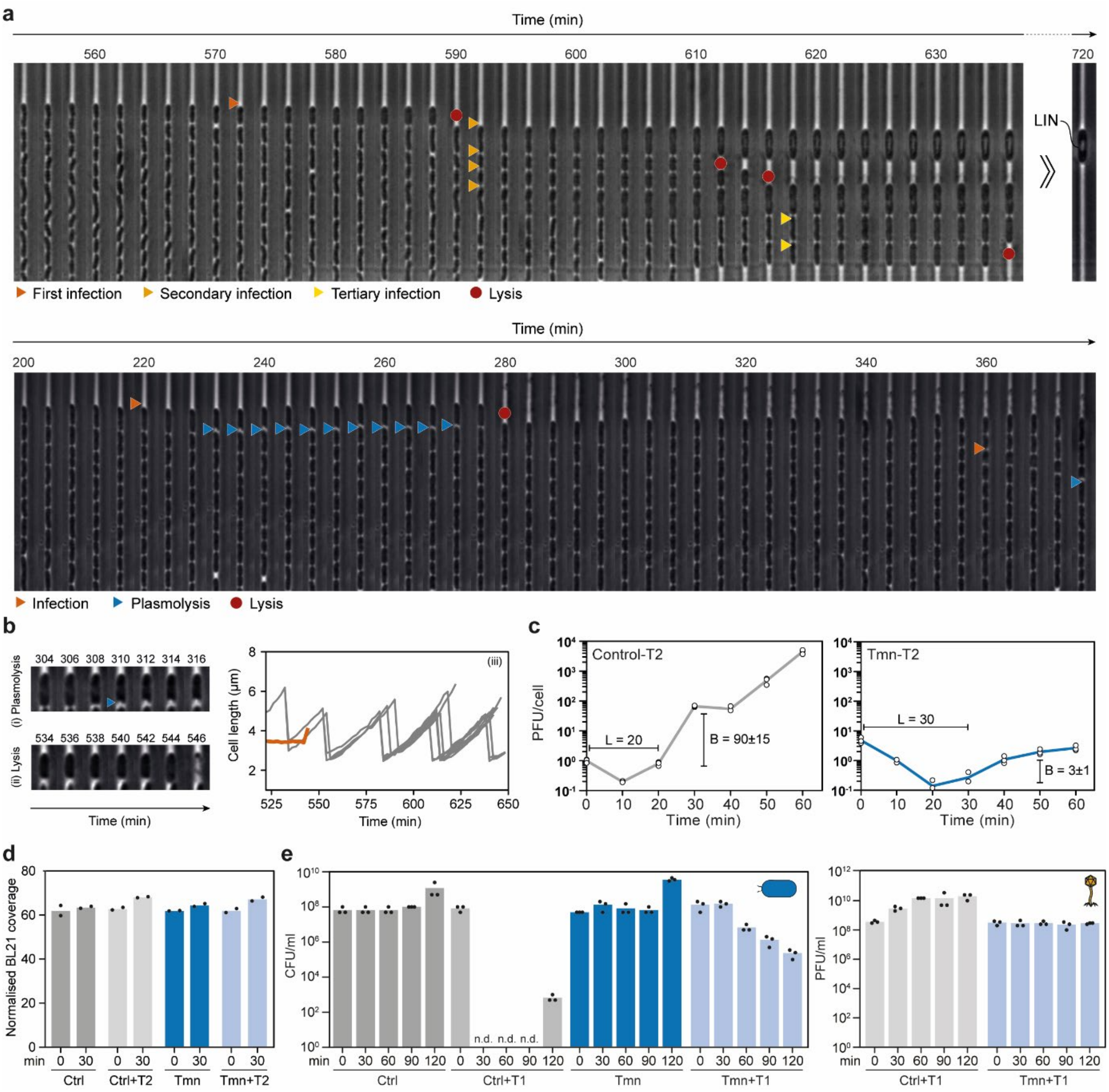


**Figure S4. Tmn-induced plasmolysis delays lysis and prevents phage transmission. a,** Time-lapse microscopy showing effective T2 infection and transmission in control cells (top) versus blocked transmission in Tmn-expressing cells (bottom). Initial infections (red) lead to secondary and tertiary infections in control cells but are halted in Tmn cells, which undergo plasmolysis (blue arrows). LIN, lysis inhibition. **b**, Zoomed examples of plasmolysis (i), lysis (ii), and successful sibling growth (iii) from Fig. 4b. **c**, One step growth curve of T2 in control (left) and Tmn-expressing (right) cells. B, burst size; L, latency period. **d**, Host genome integrity during infection measured by sequencing coverage. Bar plot shows the normalised breadth of coverage of the *E. coli* BL21 genome (fraction of genomic positions covered by mapped reads)

at 0 and 30 min for T2 infected and uninfected control (Ctrl) and Tmn-expressing cells. Dots indicate independent biological replicates, and bars show the mean. **e**, Time post infection assays measuring bacterial (CFU/ml) and phage (PFU/ml) concentrations for T1 infected and uninfected control and Tmn cells. n.d., not detectable.

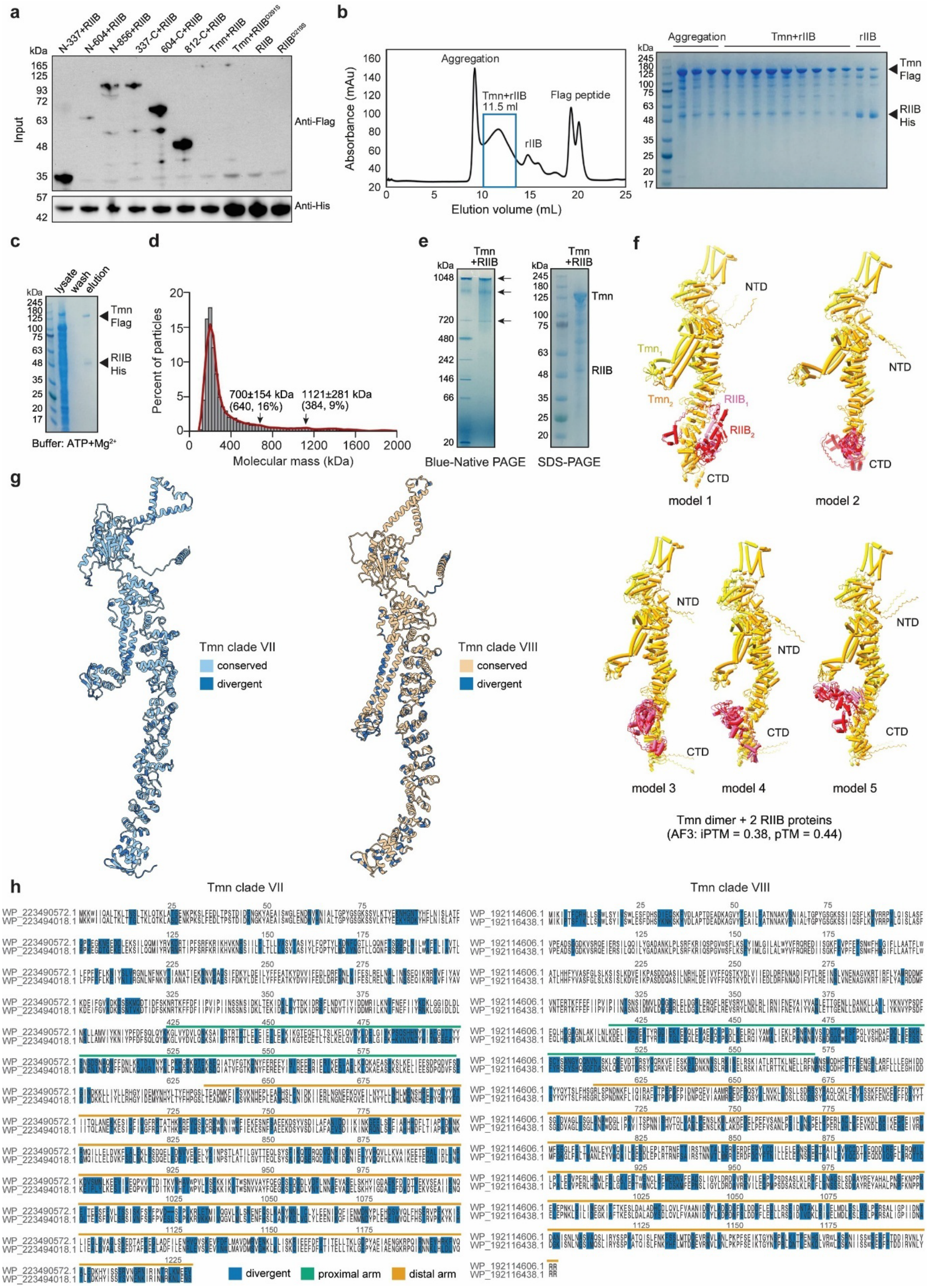


**Figure S5. Biochemical characterisation of the Tmn-RIIB interaction and RIIB variants.**

**a**, Input immunoblots corresponding to the co-IP experiments in Fig. 5d, probed with anti-Flag

(Tmn constructs) and anti-His (RIIB), indicating relative expression levels across constructs and conditions. **b,** Size exclusion chromatography (SEC) of co-expressed Tmn-25 and RIIB in *E. coli* C41 cells. Right: SDS-PAGE of SEC fractions. **c**, SDS-PAGE analysis of Tmn-Flag and RIIB-His co-expressed in *E. coli* C41 cells in the presence of 5 mM ATP and 10 mM $Mg^{2+}$. **d**, Mass photometry of purified Tmn-RIIB samples showing heterogeneous particle populations, with major distribution peaks annotated on the plot. The initial large peak is buffer background. **e**, Blue native PAGE and SDS-PAGE analysis illustrating heterogeneous assemblies and co-recovery of both proteins under these conditions. **f**, AlphaFold 3 (AF3) co-folding predictions for a Tmn-25 dimer with two RIIB monomers (five independent models shown). Confidence metrics (ipTM and pTM) are indicated. **g**, Structural comparison of two pairs of closely related Tmn homologues encoded in the same genome, highlighting clusters of substitutions mapped onto AF3-predicted structures. Left, comparison of two clade VII Tmn variants from *Sutcliffiella horikoshii* (GCF_019931755.1), with residues that differ between the proteins highlighted in dark blue on the AF3 predicted structure of WP_223490572.1. Right, comparison of two clade VIII Tmn variants from *Shewanella sp.* BF02_Schw (GCF_014596695.2), with residues that differ between the proteins highlighted in dark blue on the AF3 predicted structure of WP_192114606.1. **h**, Pairwise sequence alignment of the Tmn homologue pairs shown in (g), with residues that differ between the two variants highlighted in blue.

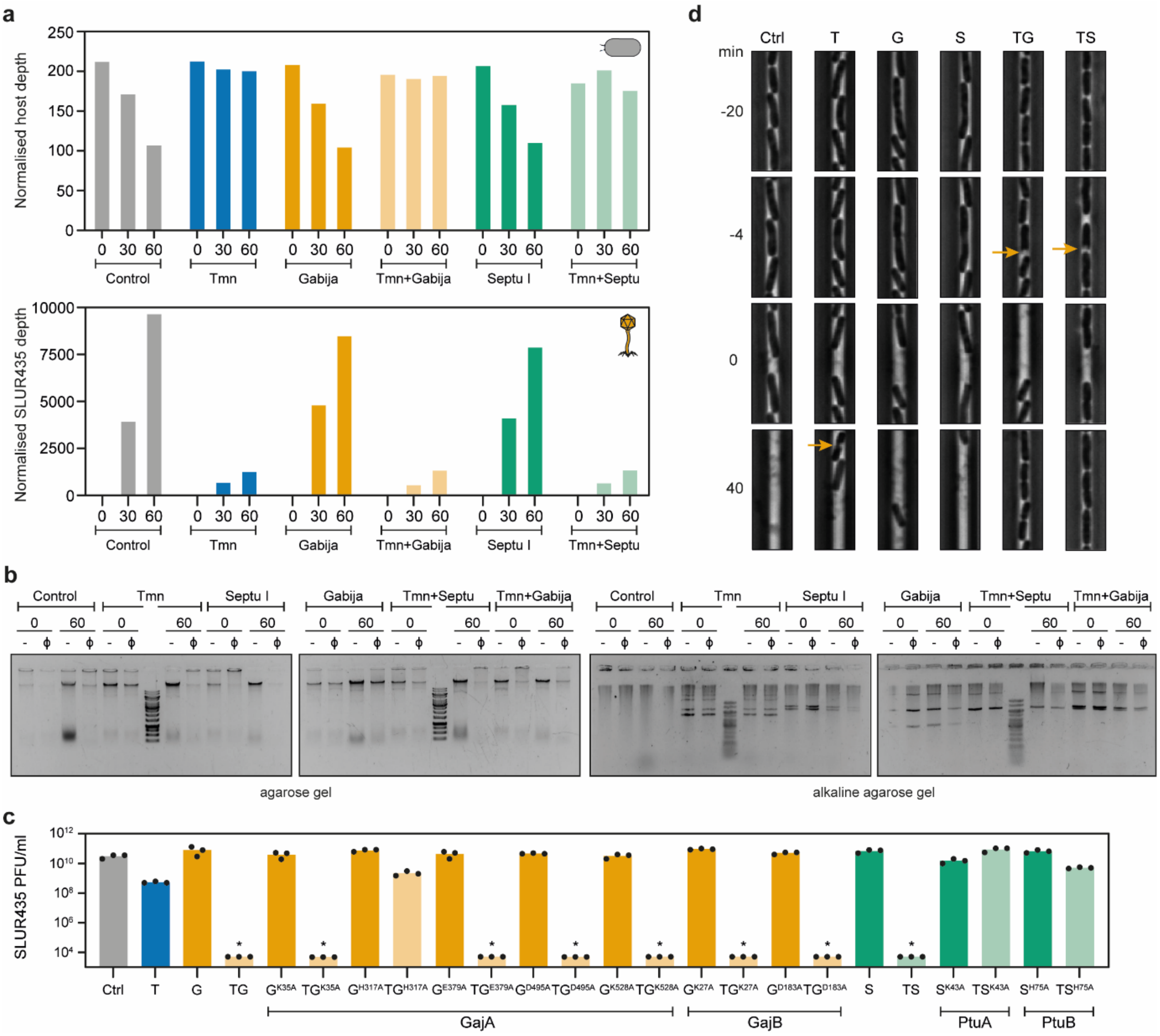


**Figure S6. Genetic and single-cell dissection of Tmn synergy with Gabija and Septu type I. a,** Quantification of bacterial host (top) and phage SLUR435 (bottom) DNA abundance during infection by Illumina sequencing. Coverage (depth) values are shown. **b**, Native (left) and alkaline (right) agarose analysis of DNA integrity during SLUR435 infection in strains expressing Tmn alone or in combination with Gabija/Septu. **c**, Mutational analysis of synergy between Tmn (T) and Gabija (G) or Septu (S) against phage SLUR435, measured as endpoint phage titre (PFU/ml). Dots indicate biological replicates; bars indicate mean. Asterisks denote significant differences ($p < 0.05$) relative to Tmn. **d,** Representative microfluidic trench trajectories illustrating infection-associated plasmolysis across strains. Example trenches are shown for control (Ctrl), Tmn (T), Gabija (G), Septu type I (S), Tmn+Gabija (TG), and

Tmn+Septu (TS). Time is indicated relative to the first detectable lysis event (0 min). Arrows indicate plasmolysis events.
